# Change in functional trait diversity mediates the effects of nutrient addition on grassland stability

**DOI:** 10.1101/2022.09.16.508233

**Authors:** Qingqing Chen, Shaopeng Wang, Eric W. Seabloom, Forest Isbell, Elizabeth T. Borer, Jonathan D. Bakker, Siddharth Bharath, Christiane Roscher, Pablo Luis Peri, Sally A. Power, Ian Donohue, Carly Stevens, Anne Ebeling, Carla Nogueira, Maria C Caldeira, Andrew S. MacDougall, Joslin L. Moore, Sumanta Bagchi, Anke Jentsch, Michelle Tedder, Kevin Kirkman, Juan Alberti, Yann Hautier

## Abstract

1. Nutrient enrichment impacts grassland plant diversity such as species richness, functional trait composition and diversity, but whether and how these changes affect ecosystem stability in the face of increasing climate extremes remains largely unknown.
2. We quantified the direct and diversity-mediated effects of nutrient addition (by nitrogen, phosphorus, and potassium) on the stability of aboveground biomass production in 10 long-term grassland experimental sites. We measured five facets of stability as the temporal invariability, resistance during and recovery after extreme dry and wet growing seasons.
3. Leaf traits (leaf carbon, nitrogen, phosphorus, potassium, and specific leaf area) were measured under ambient and nutrient addition conditions in the field and were used to construct the leaf economic spectrum (LES). We calculated functional trait composition and diversity of LES and of single leaf traits. We quantified the contribution of intraspecific trait shifts and species replacement to change in functional trait composition as responses to nutrient addition and its implications for ecosystem stability.
4. Nutrient addition decreased functional trait diversity and drove grassland communities to the faster end of the LES primarily through intraspecific trait shifts, suggesting that intraspecific trait shifts should be included for accurately predicting ecosystem stability. Moreover, the change in functional trait diversity of the LES in turn influenced different facets of stability. That said, these diversity-mediated effects were overall weak and/or overwhelmed by the direct effects of nutrient addition on stability. As a result, nutrient addition did not strongly impact any of the stability facets. These results were generally consistent using individual leaf traits but the dominant pathways differed. Importantly, major influencing pathways differed using average trait values extracted from global trait databases (e.g. TRY).
5. Synthesis. Investigating changes in multiple facets of plant diversity and their impacts on multidimensional stability under global changes such as nutrient enrichment can improve our understanding of the processes and mechanisms maintaining ecosystem stability.

## Introduction

The Earth is undergoing multiple global changes such as nutrient enrichment and climate extremes, which threaten both the diversity and stability of ecosystems (IPCC, 2023). For instance, agricultural fertilisation and atmospheric nutrient deposition have led to increased availability and redistribution of soil nutrients such as nitrogen (N), phosphorous (P), and potassium (K) globally (Galloway et al., 2021; Sardans & Peñuelas, 2015; Yuan et al., 2018). Meanwhile, climate extremes are increasing in both intensity and frequency (IPCC, 2023). Mounting evidence shows that these global changes can reduce ecosystem stability via increasing community fluctuations or indirectly via decreasing diversity (Chen et al., 2022; Hautier et al., 2015; Xu et al., 2022). However, our understanding of ecosystem stability is limited because diversity and stability are both multifaceted concepts, yet most studies only analysed one or a few facets in isolation (Chase et al., 2018; Donohue et al., 2013; Kéfi et al., 2019).

Stability characterises ecosystem responses to different types of perturbations (Pimm, 1984). In the context of climate extremes, stability of an ecosystem function (e.g., aboveground biomass production) can be defined as temporal invariability, resistance during and recovery after climate extremes (Isbell et al., 2015; Pimm, 1984). Temporal invariability indicates the degree of fluctuation and is often quantified as the ratio of the temporal mean of aboveground biomass to its standard deviation (Pimm, 1984; Tilman, 1996). While this measure is commonly termed as temporal stability in the literature, here we use temporal invariability to avoid confusion because all stability facets we investigate involve temporal dynamics. To enable comparison among sites with varying biotic and abiotic factors, resistance can be quantified as the inverse of the proportional deviation of aboveground biomass during a climate extreme from the normal level (Isbell et al., 2015). Recovery can be quantified as the proportional deviation from a normal level during a climate extreme to that after the climate extreme. Here, a normal level refers to the mean value of aboveground biomass during non-climate extremes (Isbell et al., 2015). As both resistance and recovery maintain a function around its normal level, increased resistance and/or recovery may increase temporal invariability (Isbell et al., 2015; Ives & Carpenter, 2007).

Similarly, plant diversity can be quantified in multiple dimensions, for instance, species richness, functional trait diversity, and functional trait composition (Bazzichetto et al., 2024; Craven et al., 2018). Different facets of diversity have been shown to respond differently to global changes and have different effects on ecosystem stability (Bazzichetto et al., 2024; Chen et al., 2022; Pichon et al., 2022; Suonan et al., 2023). Disentangling the direct and diversity-mediated indirect effects of global changes on multiple facets of stability is essential to understand processes and mechanisms maintaining ecosystem stability.

Past studies have highlighted the role of functional trait composition and diversity of the leaf economic spectrum (hereafter LES) in ecosystem stability (Craven et al., 2018; de Bello et al., 2021; Reich, 2014). The LES framework integrates leaf morphological, physiological, and chemical traits related to carbon acquisition and use to locate plant species along a spectrum that ranges from slow (conservative) to fast (acquisitive) strategies. Fast species can take up resources more rapidly and are typically associated with high leaf nutrients (e.g. N, P, K) and specific leaf area (Reich, 2014; Wright et al., 2004). These fast species may take advantage of increased pulses of resources and therefore recover faster after climate extremes (Bazzichetto et al., 2024; Craven et al., 2018). In contrast, slow species invest more in cell walls and secondary metabolites, having lower rates of photosynthesis and respiration (Reich, 2014; Wright et al., 2004). These features may help them endure unfavourable environments such that they may have higher resistance during climate extremes (Oram et al., 2020; Reich, 2014; Wright et al., 2004). While the ecological significance of leaf N and P has been well documented, other key elements remain less investigated (Kaspari, 2021). In particular, K is essential for regulating stomata that control gas exchange and water vapour release as well as activating enzymes for photosynthesis and protein synthesis (Kaspari, 2021). Previous single-site experiments show that nutrient addition may promote fast communities through either increasing dominance of fast species and/or shifting intraspecific traits towards fast strategies (Lepš et al., 2011; Pichon et al., 2022; Siefert & Ritchie, 2016; Tatarko & Knops, 2018; Zhou et al., 2018). However, few ecological studies on stability have as yet accounted for intraspecific trait variation, likely because of the extreme effort required to measure plant traits repeatedly. Some previous studies found that plant traits have limited explanatory power for ecosystem functioning, processes, and stability (Craven et al., 2018; van der Plas et al., 2020). These studies used species trait values from global databases (e.g. TRY) or measured in other growing environments assuming one species has a fixed trait value. Accounting for intraspecific trait shifts is important to disentangle the processes driving changes in functional trait composition and may improve prediction for ecosystem functions and stability.

Here, we use ten long-term (ranging from 10 to 15 years) standardised nutrient addition experiments to investigate the direct and diversity-mediated effects of nutrient enrichment on multidimensional ecosystem stability. We focus on five stability facets including temporal invariability of aboveground plant biomass, its resistance during and recovery after dry and wet climate extremes aggregated across a growing season (hereafter dry and wet growing seasons). We define an extreme growing season as the event occurs once per decade. We use three diversity measures including functional trait composition and diversity and species richness. We use five morphological and chemical leaf traits measured in the field accounting for intraspecific trait shifts to construct LES and calculate functional trait composition and diversity of LES and single leaf traits. To facilitate comparison with previous studies, we also use traits extracted from global trait databases.

We hypothesize that nutrient addition decreases resistance during dry and wet growing seasons. This is because nutrient addition often increases aboveground biomass, resulting in a higher normal level (Chen et al., 2023). Higher normal levels of biomass can lead to larger deviations during extreme growing seasons (e.g., decrease under dry and increase under wet growing seasons) that may exceed the nutrient-induced biomass increase in normal levels (Chen et al., 2023). During dry growing seasons, reduced water availability may reduce uptake of soluble nutrients by plants. Meanwhile, nutrient addition often increases leaf N that promotes photosynthesis and growth, which in turn increases water demands (Harpole et al., 2007). This reduced supply and increased demand of water may lead to a larger decrease in biomass (relative to the normal level) under nutrient addition than the control treatment. During wet growing seasons, increased nutrients and water availability may lead to a larger increase in biomass (relative to the normal level) under nutrient addition than the control treatment (Chen et al., 2023). Nutrient addition may also decrease recovery after dry growing seasons because increased plant mortality (due to increased normal level) can increase litter accumulation and thereby limit species colonisation (Meng et al., 2021; Southon et al., 2012). Nutrient addition may decrease recovery after wet growing seasons because increased aboveground biomass due to increased nutrients and water availability may persist or even amplify in later years (Sala et al., 2012; Wheeler et al., 2021). Moreover, nutrient addition may indirectly decrease resistance during, but increase recovery after, dry and wet growing seasons by promoting fast communities (Bazzichetto et al., 2024; Craven et al., 2018). The impact of nutrient addition on temporal invariability can, however, be weak because nutrient-induced fast communities have opposing effects on resistance and recovery (Craven et al., 2018). Furthermore, nutrient addition may indirectly decrease temporal invariability, resistance during and recovery from dry and wet growing seasons by decreasing the diversity of LES and species richness (Fig.1F). Species richness may capture diversity in phylogenetically conserved traits (e.g. plants associated with nitrogen fixation bacteria) and not conserved traits (e.g. root and size-related traits) that cannot be captured by LES. Communities with higher diversity in LES or species richness are more likely to include species that are better adapted to climate extremes. Thus, population decreases in some species may be compensated by increases in others during and after extreme growing seasons (Loreau & de Mazancourt, 2013). This leads to less aboveground biomass deviation from normal levels, which increases resistance and recovery (Bazzichetto et al., 2024) as well as temporal invariability (Craven et al., 2018). Overall, we hypothesise that nutrient addition decreases all these stability facets and that such effects are primarily mediated by changes in functional trait composition and diversity of the LES.

**Fig. 1.**
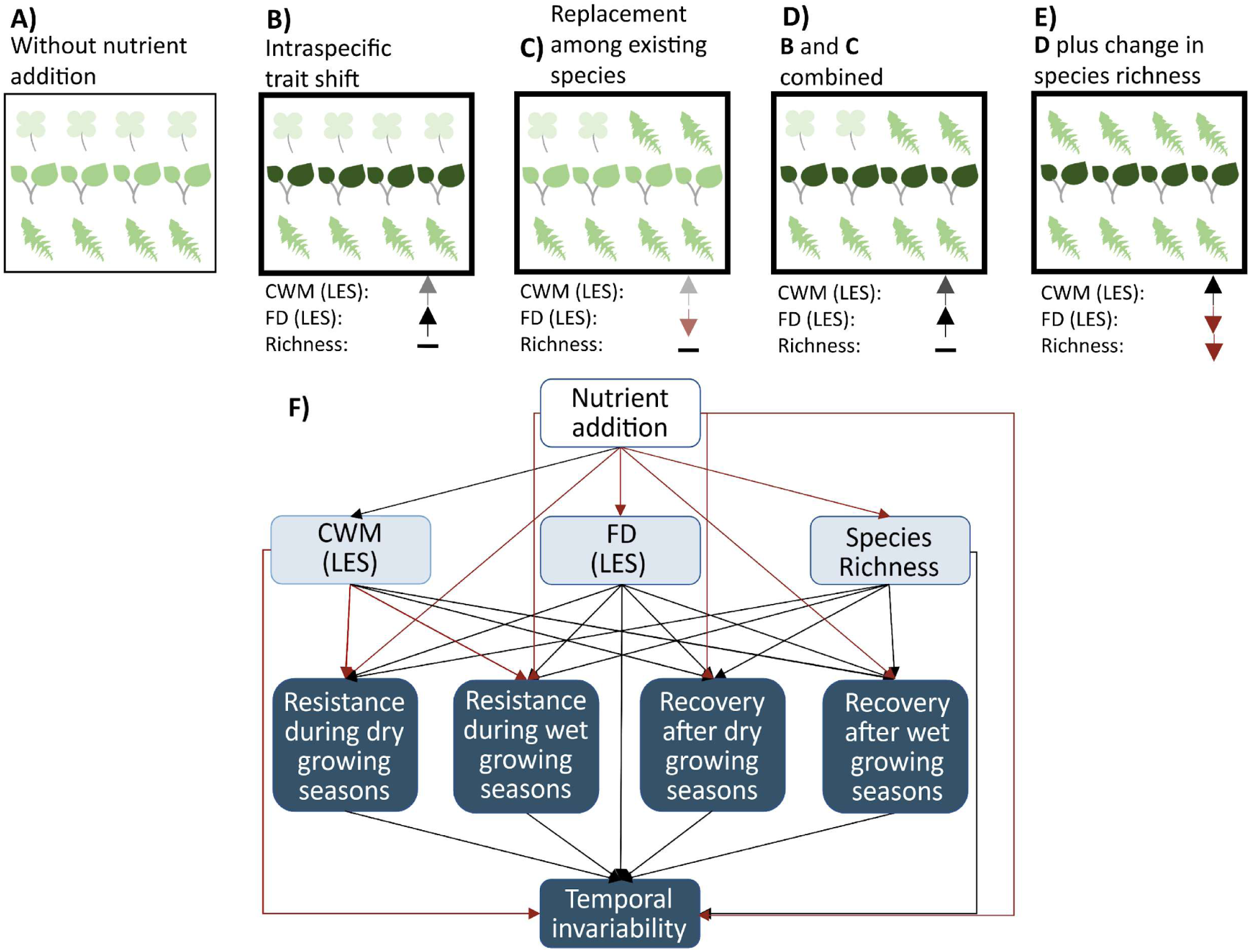
Conceptual framework illustrating how nutrient addition impacts different facets of plant diversity (A-E) and stability through its direct and diversity-mediated indirect effects (F). Darker colour in A-E represents faster species. Thick boxes represent nutrient addition conditions. Black and red arrows represent increase and decrease (darker colour represents a larger change). CWM (LES): functional trait composition as measured by community-weighted mean of leaf economics spectrum (LES); FD (LES): functional diversity of LES. Line colour in F represents positive (black) and negative (red) effects. See Table 1 for the calculation and interpretation of each variable.

**Table 1.** Variables used in this study with their mathematical definitions and interpretations. We used leaf economic spectrum (LES) as an example, we also quantified community-weighted mean (CWM) and functional diversity (FD) of single leaf traits that were used to construct LES.

| Variables | Methods | Parameters explained | Interpretations | References |
| --- | --- | --- | --- | --- |
| CWM (LES) | $\sum_{i=1}^s a_i \cdot x_i$ | $a_i$ is the relative cover of species $i$, $x_i$ is LES for species $i$. $S$ is the number of species. | A higher value indicates that a plant community is dominated by fast-growing species. | Garnier *et al.* 2004) |
| CWM induced by intraspecific trait shift | $CWM_{Nut} - CWM_{Nut*}$ | $CWM_{Nut}$ and $CWM_{control}$ are CWM in the nutrient addition and control treatment. $CWM_{Nut*}$ is the CWM in the nutrient addition treatment recalculated using LES in the control treatment. | A higher value of replacement indicates that a change in CWM is more induced by a change in relative cover among existing species while less induced by intraspecific trait shift. | (Jung *et al.* 2014) |
| CWM induced by replacement among existing species | $CWM_{Nut*} - CWM_{control}$ | | | |
| FD (LES) | $\sum_{i=1}^s a_i \cdot z_i$ | $a_i$ is cover of species $i$, $z_i$ is the distance of LES of species $i$ to community-weighted mean trait (CWM). $S$ is the number of species. | A higher value indicates that a plant community has higher diversity in LES. | (Laliberte & Legendre 2010) |
| Temporal invariability | $u/\sigma$ | $\mu$ is the mean of aboveground biomass, $\sigma$ is the standard deviation of it over time. | A higher value indicates that a plant community fluctuates less in aboveground biomass over time. | (Pimm 1984) |
| Resistance during dry/wet growing seasons | $\frac{\bar{Y}_n}{\vert Y_e - \bar{Y}_n\vert }$ | $\bar{Y}_n$, $Y_e$, and $Y_{e+1}$ are aboveground biomass during normal growing seasons, during a dry or wet growing season, and one year after a dry or wet growing season, respectively. | A higher value indicates that aboveground biomass deviates less during a dry or wet growing season from that of the average of normal growing seasons. | (Isbell *et al.* 2015) |
| Recovery after dry/wet growing seasons | $\frac{\vert Y_e - \bar{Y}_n\vert }{\vert Y_{e+1} - \bar{Y}_n\vert }$ | | A higher value indicates that aboveground biomass deviates less one year after a dry or wet growing season relative to that during a dry or wet growing season. | (Isbell *et al.* 2015) |

## Materials and Methods

### Experimental Design

We used a coordinated, multisite, and multiyear nutrient addition and herbivore manipulation experiment (NutNet; www.nutnet.org) initiated in 2007 (Borer et al., 2014, 2017). The original design includes a factorial manipulation of nutrients (N, P & K) plus two fences to exclude herbivores (one without nutrients addition and the other with NPK). Each treatment was imposed to a 25 m^2^ plot and replicated in at least three blocks. N was supplied as slow-release urea ((NH_2_)_2_CO), P was supplied as triple superphosphate (Ca(H_2_PO_4_)_2_), and K as potassium sulphate (K_2_SO_4_). N, P, and K were added annually at a rate of 10 g m^-2^ y^-1^ (i.e 100 kg/ha/year). A micronutrient and macronutrient mix (Fe, S, Mg, Mn, Cu, Zn, B, and Mo in combination) was applied at a rate of 100 g m^-2^ once at the start of the experiment, as part of the K addition. Further details on the design are available in (Borer et al., 2014). No permits were required for fieldwork. We use traits measured in all treatments (see section ‘Leaf trait measurements’ for details), we focus on exploring stability in the control and nutrient addition (NPK) treatments.

### Site selection

We selected ten long-term (ranging from 10 to 15 years) sites because they had: 1) both control and nutrient addition (NPK) treatments and the occurrence of at least one extreme dry and one extreme wet growing season during the experiment. See section ‘Defining climate extremes and stability facets’ for detail in classifying dry and wet growing seasons; 2) three blocks. For sites with more than three blocks, we used the first three blocks according to the block number recorded by site PIs; 3) more than three plant species measured for leaf traits in both control and nutrient addition conditions; 4) >50% proportional cover of species with trait values in a plant community averaged across blocks and experimental years. So, the community-weighted mean (CWM) and functional diversity (FD) of traits can reasonably represent the whole plant community. See section ‘Leaf trait measurements’ for details. These sites are distributed in North America (8 sites) and Australia (2 sites) (Fig. S1; Table S1). Data are achieved in Figshare (Chen et al. 2024). These sites are dominated by herbaceous plant species, covering montane, alpine, and semiarid grasslands as well as prairies and old fields, which we refer to as grasslands for simplicity (see Table S1 for geolocation, climate, and experimental duration for these sites).

### Sampling protocol

All sites followed standard NutNet sampling protocols. A 1×1m subplot was permanently marked within a 25 m^2^ plot. Number of species, species identity, and their covers were recorded once per year in these 1×1 m subplots at most sites. At a few sites with strong seasonality, cover and biomass were recorded twice per year to include a full list of species and follow typical management procedures. For those sites, the maximum cover for each species and total biomass for a community were used in the analysis. The taxonomy was checked and adjusted within sites to ensure consistent naming over time. For instance, when individuals could not be identified as species in all years, they were aggregated at the genus level but referred to as taxa for simplicity. Meanwhile, aboveground biomass was measured adjacent to these permanent subplots within two 1 × 0.1 m strips (in total 0.2 m^2^), which were moved from year to year to avoid resampling. All aboveground biomass was clipped, sorted into dead and live, and dried at 60 °C to constant mass before weighing to the nearest 0.01 g.

### Leaf trait measurements

Leaf morphological and chemical traits including leaf carbon, leaf nitrogen, leaf phosphorus, leaf potassium, specific leaf area (hereafter leaf C, N, P, K, SLA) were measured after 2, 3, or 4 years of nutrient addition (details in Table S2). Traits were measured for 3 to 5 of the most abundant species (ranked by cover) in each subplot according to standard protocols (Pérez-Harguindeguy et al., 2016). A detailed description of these trait measurements can be found in (Firn et al., 2019). Briefly, for each species measured for leaf traits, five fully grown leaves without clear grazing marks were randomly selected. Leaf area (mm^2^) was measured using a leaf area metre or a scanner. After that, dry weight (g) of leaves was measured after oven-drying at 60 °C for 48 h. Specific leaf area (SLA; mm^2^g^-1^) was calculated as leaf area divided by dry weight. Then, leaf nutrient concentrations (%) including C, N, P, and K were determined. Leaf P, K, and C were determined using laser ablation inductively coupled plasma mass spectrometry following (Duodu et al., 2015). Leaf N was determined using a LECO TruMac, based on a combustion technique using thermal conductivity relative to pure gas with an error < 1%.

We separated leaf traits measured in ambient (control, Fence) and nutrient-enriched conditions (N, P, K, NP, NK, PK, NPK, NPK+Fence). Additionally, multiple trait measurements for the same species at one site (e.g., from different blocks or nutrient treatments) were averaged. We did this to maximise the number of species with available trait data and because previous results found little variation in leaf traits among blocks within the ambient and nutrient-enriched conditions at 27 NutNet sites (Firn et al., 2019). Due to this aggregation, a larger number of species measured for traits and proportional cover of species with traits in plant communities were found under nutrient addition than ambient conditions at nearly all sites (Table S2). Across all sites, plant species with trait values account for >52% of the cover; at 6 of 10 sites this proportion was similar under control and nutrient addition treatments (Table S2). Overall, 102 plant species were measured for the five leaf traits and 92 species have data for all these traits.

### The leaf economic spectrum, community-weighted mean traits, and functional trait diversity

We used the five leaf traits from the 92 species to construct leaf economic spectrum (LES) using principal component analysis (PCA) as coded in the “PCA” function from the R package “FactoMineR” (Lê et al., 2008). We extracted the first axis of the PCA, which explained 34.7% of the variance, to represent the LES. Higher values (i.e. lower leaf C, higher N, P, K, and SLA) indicate faster species (Fig. S2a). Due to relatively low variance explained in the first axis of the PCA, we calculated CWM and FD of LES as well as of individual leaf traits. CWM is the cover-weighted average of each trait in a community (Garnier, et al., 2004). FD is cover-weighted dispersion of each trait relative to CWM (Laliberte & Legendre, 2010). We calculated CWM and FD using the function “dbFD” from the R package “FD” (Laliberté et al., 2014). See Table 1 for mathematical formulas for CWM and FD and their interpretations. These variables were calculated annually for each subplot and then averaged across years.

### Partitioning community-weighted mean traits into intraspecific trait shift and species replacement

In addition to calculating CWM based on all species with trait data, we recalculated it for shared species (species that were present in both the control and nutrient addition subplots). This allowed us to partition CWM based on shared species into intraspecific trait shift and replacement of existing species following (Jung et al., 2014). See Table 1 for mathematical formulas for calculating and interpreting these variables.

### Compare results using trait values extracted from global trait databases

To compare with previous studies that quantify LES based on global trait databases that often include leaf dry matter content (LDMC), we compiled species-level trait data (leaf C, N, P, K, SLA, LDMC) from TRY (Version 6; Kattge et al., 2020), BIEN (Version 1.2.6; Maitner et al., 2018), AusTraits (Version 5.0.0; Falster et al., 2021) for NutNet species (Chen 2024). Following (Craven et al., 2018), all traits were first averaged within databases and then across them for each species regardless of their geolocation. Overall, species-level traits covered less than 50% of the species occurring at these 10 sites. But species with extracted leaf N and SLA data accounted for > 50% of community cover at most NutNet sites (Table S3), thus we use these two species-level traits to calculate CWM and FD. We compared these results to those based on traits directly measured in the field. Moreover, only 31 species have data for all these six leaf traits. To increase trait coverage, following (Craven et al., 2018), missing species-level traits were filled using the average trait value from other species in the same genus for which trait values were available. To ensure that filled trait values were not biassed towards species with a higher number of records, trait values were first averaged for each species, then averaged across species within a genus. Because of low coverage of leaf K data for species from these 10 NutNet sites, here we used leaf C, leaf N, leaf P, SLA, and LDMC to construct LES. Similarly, we extracted the first axis of PCA, which explained 40.3% of the variance, to represent the LES (Fig. S2b).

### Defining climate extremes and stability facets

We used the standardised precipitation–evapotranspiration index (SPEI) to classify climate extremes for each site. SPEI was calculated as the standardised (z-score) water balance (precipitation – evapotranspiration; mm) over the growing season from 1901 to 2022. We used water balance during growing seasons because previous studies show it is better correlated with aboveground biomass than total annual water balance (Robinson et al., 2013). Growing seasons were defined by the site PIs (Table S1). Precipitation and potential evapotranspiration used to calculate SPEI were downloaded from https://crudata.uea.ac.uk/cru/data/hrg/cru_ts_4.07/ (accessed (Harris et al., 2020). Following (Isbell et al., 2015), we categorised each growing season into normal, dry, and wet using the cutoffs of 0.67 and 1.28 SD (1.28: occurring once per decade; 0.67: once every four years; SD: standard deviation). That is, normal growing season: -0.67 SD < SPEI < 0.67 SD; dry: SPEI ≤ -1.28 SD; and wet: SPEI ≥ 1.28 SD. In total, 64, 19, and 19 normal, dry, and wet growing seasons across sites were detected in our data. When two (or more) extreme growing seasons of the same kind happen consecutively (e.g., wet followed by wet), recovery was only calculated for the last growing season, which must be followed by a normal or a less extreme growing season (those between normal and extreme).

We quantified resistance as the inverse of the proportional deviation of aboveground biomass from normal levels during a dry or wet growing season. Also, we quantified recovery as the inverse of the proportional lack of recovery in aboveground biomass during the year after a dry or wet growing season following (Isbell et al., 2015). We treated resistance during and recovery after dry and wet growing seasons individually, and averaged each over experimental years to match the data structure of temporal invariability. We quantified temporal invariability as the ratio of the temporal mean to the standard deviation of aboveground biomass in each plant community (Pimm, 1984). To eliminate potential trends in aboveground biomass over time, we calculated detrended standard deviation from the residuals of a linear model (function “lm”) regressing aboveground biomass against experimental years (Tilman et al., 2006). See Table 1 for mathematical formulas for calculating stability facets and their interpretations.

### Statistical analysis

All analyses were performed in R v.4.1.6 (R Core Team., 2022). R codes are published (Chen 2024). We used linear mixed-effects models (function “lme”) from the R package “nlme” (Pinheiro et al., 2017) for the following analyses. We built models where site and block nested within the site were the random effects and treatment was the fixed effect. First, we tested whether nutrient addition impacted CWM and FD of various traits using all species with trait data and the shared species. We also tested whether nutrient addition impacted intraspecific trait shifts and species replacement (drivers for CWM) of various traits based on shared species. Second, we examined the effects of nutrient addition on each stability facet. To that end, we first disentangled how nutrient addition impacted resistance and recovery through aboveground biomass deviation under extreme growing seasons from that during normal growing seasons. We aggregated aboveground biomass and the magnitude of aboveground biomass deviation (values are positive only) from normal levels during and one year after dry and wet growing seasons, in control and nutrient addition treatments across sites. We present raw data for aboveground biomass under different growing seasons over the experimental years at each site (Fig. S3). Then, we tested whether nutrient addition impacted stability facets.

We built structural equation models (SEMs) to evaluate the direct effects of nutrient addition on stability facets as well as its indirect effects through CWM, FD, and species richness. The SEMs were built using the function “psem” from the R package piecewiseSEM (Lefcheck, 2016). An initial model was built based on prior knowledge (Fig. 1F). For each component model in SEM, we used the function “lme” with site and block nested within site as random effects. We estimated variance inflation for each component model to check whether multicollinearity affects parameter estimates, which were smaller than 2 in all component models. The goodness of fit of SEM models were assessed by Fisher’s C statistic, with a higher *p* value (e.g. ≥0.05) indicating a good model fit. We used CWM and FD of LES and each measured leaf trait in the model to link to facets of stability under nutrient addition. We also used species-level leaf N, SLA, and LES based on species- and genus-level filled traits from global trait databases to link them to facets of stability under nutrient addition.

## Results

Using all species having traits, nutrient addition decreased FD while increasing CWM of LES (Fig. 2). That is, nutrient addition led to faster communities. The result for CWM was similar using only shared species (i.e., species occurring in both control and nutrient addition treatments). Using the shared species, we further found that the increased CWM of LES under nutrient addition was driven mainly by intraspecific trait shifts rather than replacement among existing species (Fig. 2; Table S4). Nutrient addition also decreased FD of leaf C, N, P, but had no effects on FD of leaf K and SLA (Fig. 2; Table S4). Nutrient addition also increased CWM of leaf N, P, K, and SLA, but had no effect on CWM of leaf C. Using the shared species, we found changes were again mainly driven by intraspecific trait shifts, but the increased CWM of leaf P was also partly driven by species replacement among existing species (Fig. 2; Table S4).

**Fig. 2.**
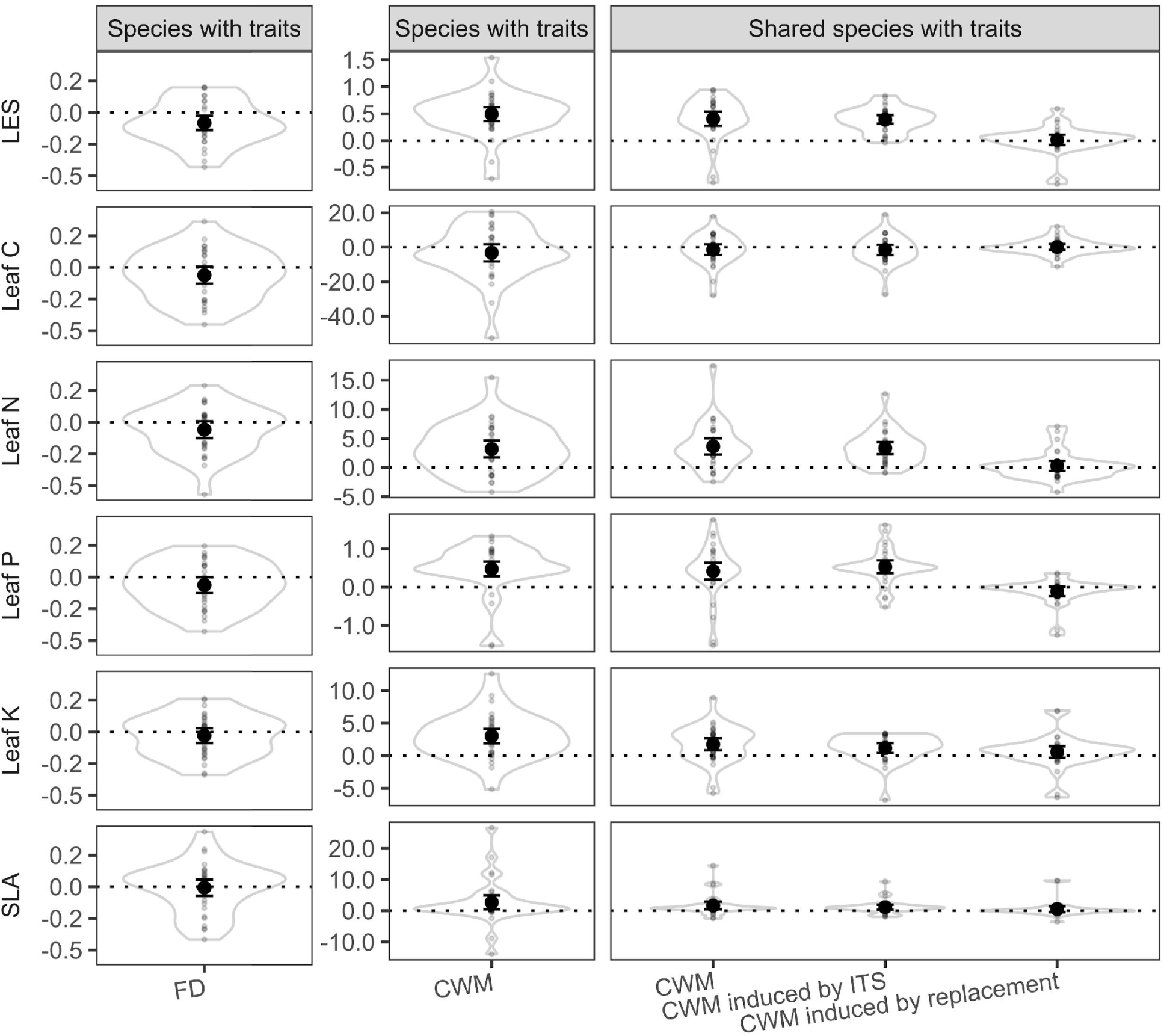
Effects of nutrient addition on functional diversity (FD) and community-weighted mean (CWM) of the leaf economic spectrum (LES) and single leaf traits used to construct LES. For shared species (i.e. species present in both control and nutrient addition treatments), change in CWM was further attributable to intraspecific trait shifts (ITS) and replacement among existing species. Leaf C, Leaf N, Leaf P, Leaf K, SLA are leaf carbon, nitrogen, phosphorus, potassium, specific leaf area, respectively. Small points are effects of nutrient addition on each community-level trait from each block at each site. Violin shapes show distribution of values. Large black points are mean values over all 10 sites estimated from linear mixed effect models, error bars are 95% confidence intervals. See Table S4 for test statistics.

During normal growing seasons, nutrient addition significantly increased aboveground biomass by 60% (Fig. 3; Table S5). During dry growing seasons, aboveground biomass decreased relative to their normal levels under both control and nutrient addition treatments, but this decrease was more pronounced under nutrient addition. Nutrient addition weakly increased aboveground biomass deviation (i.e., absolute difference between dry and normal seasons) by 9% (Table S5). One year after dry growing seasons, biomass generally returned to their normal levels. The deviation in biomass was, however, 104% higher under nutrient addition than the control, suggesting the biomass recovery was more variable (some sites increased while others decreased) under nutrient addition. During wet growing seasons, aboveground biomass increased relative to normal levels under both control and nutrient addition treatments. This increase was more pronounced under nutrient addition, with biomass deviation 68% higher under nutrient addition than the control. These deviations persisted to the year following a wet growing season (Fig. 3; Table S5).

**Fig. 3.**
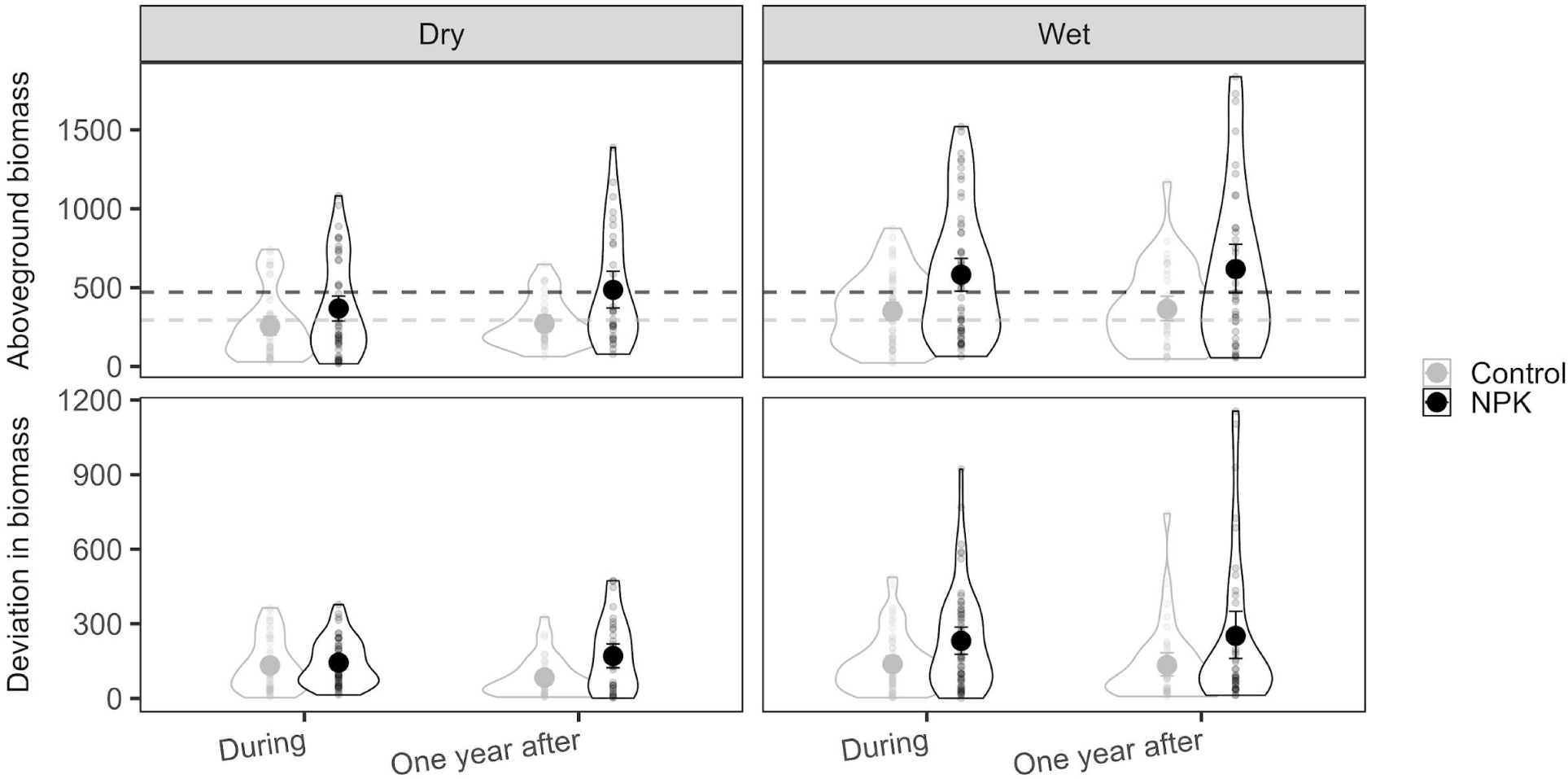
Aboveground biomass (gm^-2^) and the magnitude of its deviation (gm^-2^) during and after dry and wet growing seasons from that of normal levels. These variables were used to quantify resistance and recovery. Deviation in biomass refers to absolute change in aboveground biomass from normal levels. Dashed lines indicate mean aboveground biomass during normal growing seasons (i.e., normal levels). Note, each treatment in each block at each site had its own normal level. Small points are values from each block at each site. Violin shapes show distribution of values. Large points indicate average values across 10 sites. Error bars are 95% bootstrapped confidence intervals. See Table S5 for test statistics.

The SEM revealed that nutrient addition impacted facets of stability directly and indirectly through different facets of plant diversity (Fig. 4A; Table 2). When using CWM and FD of LES in the SEM, nutrient addition increased resistance during dry growing seasons mainly through decreasing FD of LES. Nutrient addition influenced resistance during wet growing seasons both directly and indirectly, with the positive direct effects partially offset by the negative indirect effects through reducing FD of LES. This resulted in a weak overall increase in resistance during wet growing seasons under nutrient addition. Nutrient addition decreased recovery after dry growing seasons through decreasing FD of LES, but had no effect on the recovery after wet growing seasons. Nutrient addition impacted temporal invariability weakly through the indirect effects mediated by resistance and diversity facets. Overall, these pathways resulted in weak effects of nutrient addition on all facets of stability (Fig. 4A; Table 2; Fig. S4; Table S6).

**Fig. 4.**
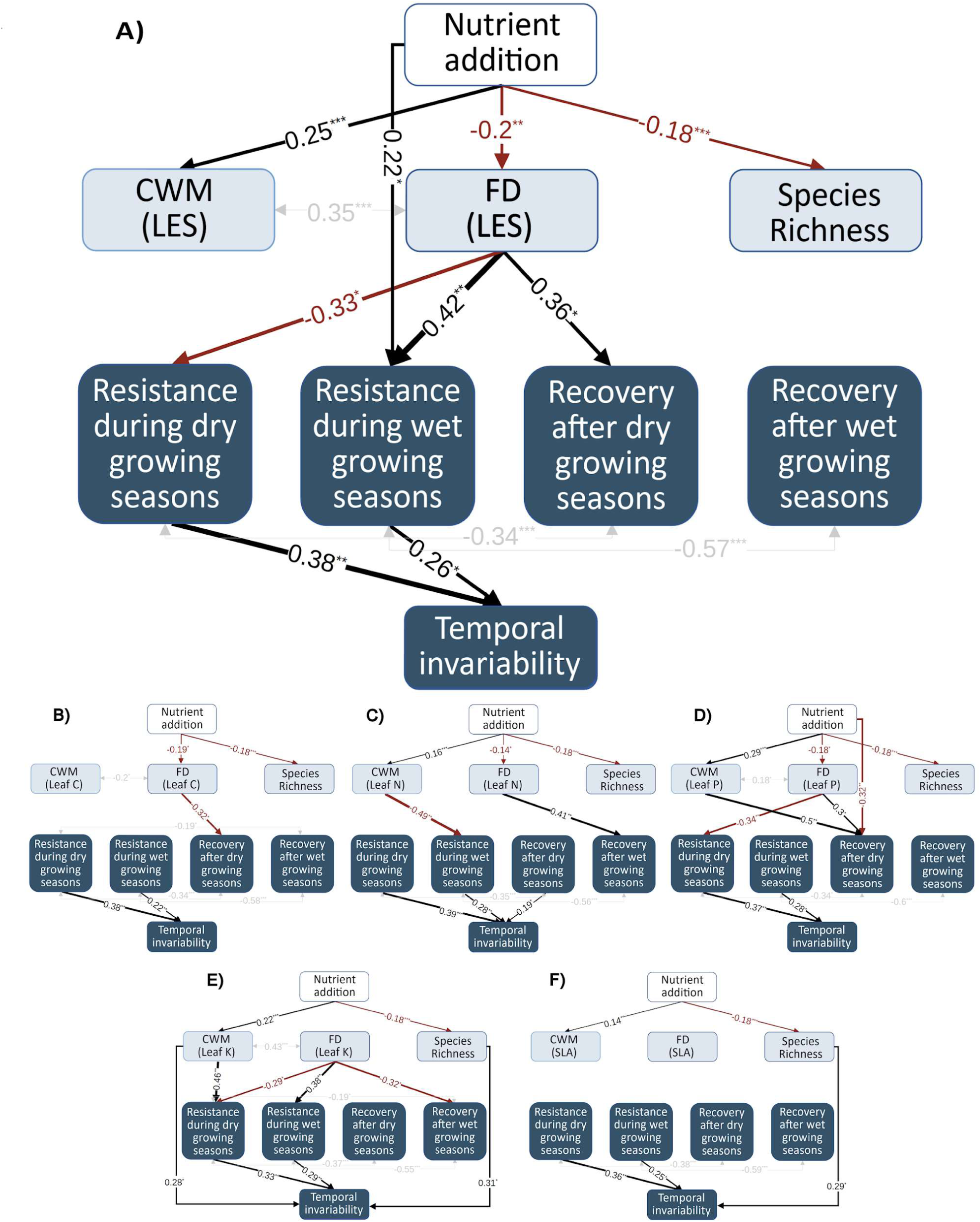
Direct and diversity-mediated effects of nutrient addition on facets of stability, where functional trait diversity and composition is based on the leaf economic spectrum (A), leaf carbon (B), leaf nitrogen (C), leaf phosphorus (D), leaf potassium (E), and specific leaf area (F). Arrows represent relationships among variables. The displayed numbers are standardised path coefficients. The width of arrows indicates the strength of the pathways. Line colour represents positive (black) and negative (red) effects. Non-significant paths are not shown. Grey double-headed arrows and text show correlated errors. Asterisks indicate significant paths: * p ≤ 0.1; ** p ≤ 0.05; *** p ≤ 0.001. All stability facets were on the log scale to improve normality and homogeneity of variance. See Table 2 for the breakdown of direct and diversity-mediated indirect effects of nutrient addition on facets of stability. All models fit the data well, see Table S7 for goodness of model fit and variance explained (R^2^) for each component model.

**Table 2.** Direct and diversity-mediated effects (standardised path coefficients) of nutrient addition on facets of stability. Results were summarised from both significant and non-significant paths from structure equation models in Fig. 4. LES, Leaf C, Leaf N, Leaf P, Leaf K, SLA are the leaf economic spectrum, leaf carbon, nitrogen, phosphorus, potassium, specific leaf area, respectively.

| Trait name | Effects/pathways | Resistance during dry growing seasons | Resistance during wet growing seasons | Recovery after dry growing seasons | Recovery after wet growing seasons | Temporal invariability |
| --- | --- | --- | --- | --- | --- | --- |
| LES | Direct effects | 0 | 0.22 | -0.17 | -0.18 | -0.1 |
|  | Direct effects through resistance and recovery |  |  |  |  | 0.01 |
|  | Indirect effects through diversity facets | 0.16 | -0.14 | -0.01 | 0.05 | -0.02 |
|  | Indirect effects through resistance and recovery (and through diversity facets) |  |  |  |  | 0.03 |
|  | Total effects | 0.16 | 0.08 | -0.18 | -0.13 | -0.09 |
| Leaf C | Direct effects | 0.1 | 0.13 | -0.21 | -0.12 | -0.05 |
|  | Direct effects through resistance and recovery |  |  |  |  | 0.01 |
|  | Indirect effects through diversity facets | 0.05 | -0.05 | 0.03 | -0.01 | -0.06 |
|  | Indirect effects through resistance and recovery (and through diversity facets) |  |  |  |  | 0.01 |
|  | Total effects | 0.16 | 0.08 | -0.18 | -0.13 | -0.09 |
| Leaf N | Direct effects | 0.1 | 0.15 | -0.17 | -0.13 | -0.09 |
|  | Direct effects through resistance and recovery |  |  |  |  | 0.03 |
|  | Indirect effects through diversity facets | 0.05 | -0.07 | -0.01 | 0 | -0.03 |
|  | Indirect effects through resistance and recovery (and through diversity facets) |  |  |  |  | 0 |
|  | Total effects | 0.16 | 0.08 | -0.18 | -0.13 | -0.09 |
| Leaf P | Direct effects | 0.01 | 0.15 | -0.32 | -0.16 | -0.11 |
|  | Direct effects through resistance and recovery |  |  |  |  | -0.04 |
|  | Indirect effects through diversity facets | 0.14 | -0.07 | 0.14 | 0.03 | 0 |
|  | Indirect effects through resistance and recovery (and through diversity facets) |  |  |  |  | 0.06 |
|  | Total effects | 0.16 | 0.08 | -0.18 | -0.13 | -0.09 |
| Leaf K | Direct effects | 0.02 | 0.12 | -0.2 | -0.14 | -0.12 |
|  | Direct effects through resistance and recovery |  |  |  |  | -0.01 |
|  | Indirect effects through diversity facets | 0.14 | -0.04 | 0.02 | 0.01 | 0.01 |
|  | Indirect effects through resistance and recovery (and through diversity facets) |  |  |  |  | 0.04 |
|  | Total effects | 0.16 | 0.08 | -0.18 | -0.13 | -0.09 |
| SLA | Direct effects | 0.1 | 0.1 | -0.21 | -0.12 | -0.07 |
|  | Direct effects through resistance and recovery |  |  |  |  | 0.01 |
|  | Indirect effects through diversity facets | 0.05 | -0.02 | 0.03 | -0.01 | -0.04 |
|  | Indirect effects through resistance and recovery (and through diversity facets) |  |  |  |  | 0.02 |
|  | Total effects | 0.16 | 0.08 | -0.18 | -0.13 | -0.09 |

Similar results were revealed when using CWM and FD of single leaf traits, although the dominant influencing pathways differed (Fig. 4B-4F; Table 2). Also, the diversity-mediated indirect pathways were stronger using leaf N, leaf P, and leaf K, than leaf C, and SLA. For instance, nutrient addition did not strongly impact any stability facet through FD of SLA. When using CWM and FD of traits extracted from global databases (Fig. S5; Fig. S6; Fig. S7; Table S7; Table S8), nutrient addition did not alter CWM whereas it had a strong negative effect on temporal invariability through decreasing species richness. When including CWM and FD of leaf N and SLA based on species-level traits extracted from global trait databases, nutrient addition had strong effects on some stability facets through FD (Fig. S6; Fig. S7; Table S7; Table S8).

## Discussion

Our study tested the role of the leaf economic spectrum (LES) and single leaf traits by accounting for intraspecific trait variations in mediating ecosystem stability following nutrient addition. Using leaf traits measured in the field, we quantified the contribution of intraspecific trait shifts and species replacement to change in functional trait composition as responses to nutrient addition and its implications for ecosystem stability. Our study also expanded the scope of stability analyses by including five facets (resistance during and recovery after dry and wet growing seasons, and temporal invariability). Among these, resistance during and recovery after wet growing seasons have been largely overlooked by previous studies. We found that nutrient addition strongly impacted functional trait composition and diversity of LES and single leaf traits, and change in functional trait composition was mainly driven by intraspecific trait shifts. This suggests that intraspecific trait shifts need to be included for accurately predicting ecosystem stability under global changes such as nutrient enrichment. The changes in plant diversity mediated changes in different facets of ecosystem stability under nutrient addition. The major influencing pathways differed using traits measured on-site and that extracted from global databases.

### The role of intraspecific trait shifts in community trait change

Nutrient addition promoted fast communities while decreasing FD of LES through intraspecific trait shifts and changing species richness (Fig. 1E). The change in CWM of LES was mainly induced by intraspecific trait shifts corroborates results from previous studies under other global change scenarios (Jung et al., 2014; Duodu et al., 2015; Niu et al., 2016; Tatarko & Knops, 2018; Oram et al., 2020). These results indicate that plant species may rapidly adapt/acclimate to global change factors including flooding, drought, and nutrient enrichment. At many NutNet sites, leaf traits were measured after 4 years of nutrient addition, possibly resulting in higher intraspecific trait shifts. Although lacking a test, species replacement that takes longer to manifest may become more important than intraspecific trait shifts in determining community traits in the longer term. A previous NutNet study shows that leaf nutrients, but not SLA, of abundant species are consistent indicators of increased soil nutrients (Firn et al., 2019). Extending that, our results suggest community-level leaf P and leaf N were more responsive than leaf K, leaf C, and SLA to nutrient addition.

### Diversity-mediated effects on facets of stability

Plant traits have long been viewed as a common currency to link ecosystem processes, functions, and stability (de Bello et al., 2021; Funk et al., 2017). Past studies have shown that FD and CWM of LES and single leaf traits can be tightly linked to temporal invariability (Craven et al., 2018; Schnabel et al., 2021; Suonan et al., 2023), resistance during and recovery after dry and wet climate extremes (Bazzichetto et al., 2024; Oram et al., 2020). For instance, using bivariate analysis, (Oram et al., 2020) found that CWM, but not FD, of LES was strongly related to resistance during and recovery from floods. (Mariotte et al., 2013) also found that subordinate plant species enhance community resistance during a summer drought in a semi-natural grassland. However, (Bazzichetto et al., 2024) found that FD of LES was positively related to drought resistance, but CWM of LES was positively related to recovery after short-term droughts (aggregated over 3 months prior to biomass harvest). Surprisingly, the commonly-assumed trade-off between CWM of LES (i.e. slow and fast communities) on resistance and recovery was not supported in these studies as well as not in ours (Bazzichetto et al., 2024; Craven et al., 2018; Oram et al., 2020). We found that FD of LES was better linked to stability than CWM of LES and species richness under nutrient addition (Fig. 4A), suggesting that compensatory dynamics among species may be more important than dominant species in driving community stability at these sites. Interestingly, we found that nutrient addition had opposing effects on resistance during dry and wet growing seasons through decreasing FD of LES (Fig. 4A). That is, plant communities with higher functional diversity of LES exhibited higher resistance (smaller biomass deviation) than those with less diverse LES during wet growing seasons, likely attributed to compensatory dynamics. The opposite effects during dry growing seasons may be because more diverse communities lost more biomass. As different traits respond differently to climate extremes, they could mediate ecosystem functions and stability in different ways (de Bello et al., 2021; Funk et al., 2017). Indeed, we found that results based on LES differed from those based on single leaf traits in dominant pathways. Results also differ among single leaf traits with leaf P and leaf N showing stronger links.

That said, the effects of nutrient addition on stability mediated by functional trait composition and diversity were relatively weak. This may be due to the following reasons. First, although the LES is a well-established concept and its potential connection to ecosystem stability is well described (de Bello et al., 2021), the link between LES and biomass production is only suggestive. Indeed, from a physiological perspective, plants could increase aboveground biomass by structural elements (e.g. increasing the number of leaves, stems) instead of increasing SLA and leaf nutrients (Firn et al., 2019). Second, we may miss some potentially important traits such as root size or metabolic traits (Bazzichetto et al., 2024; Schnabel et al., 2021). Root traits are particularly important for plant species to access soil water and nutrients. However, a greenhouse experiment found that the LES can better predict biomass resistance during and recovery from flooding than the root economic spectrum (Oram et al., 2020). So far, a lack of studies focusing on root and metabolic traits limits our ability to draw any solid conclusions (Oram et al., 2020; Schnabel et al., 2021), and future work resolving the role of such traits could offer new insights into the trait-stability framework. Third, we did not measure traits for all species and those rarer species may strengthen or alter the results found here (Sun et al., 2022). Fourth, we did not consider intraspecific trait shifts over time, which can be a major source of community-level trait shifts (Wheeler et al., 2022), and further impact the prediction of community-level traits on ecosystem stability. This temporal intraspecific trait shift is ignored in many ecological studies using traits to predict ecosystem functions and stability, likely because of the extreme effort required. Last but not least, as our study focused on patterns across sites, we partially accounted for the heterogeneity of climate conditions at each site using site-specific normal levels and extreme growing seasons. Considering environmental dependence together with trait responses, and ecosystem functions, processes, and stability simultaneously would improve our understanding of their links and underlying mechanisms.

### Comparison between analyses using trait values from different sources

To our knowledge, no studies so far have compared the effects of community-level traits measured in the field accounting for intraspecific trait variation with those extracted from global trait databases that often do not account for intraspecific trait variation on ecosystem stability. Increasing accumulation of measured plant traits globally (e.g. TRY database) have greatly advanced the field of trait-based studies (Kattge et al., 2020). Previous studies using trait values extracted from global trait databases to explain ecosystem functions and stability often assume one species has one fixed trait value (Bazzichetto et al., 2024; Craven et al., 2018; van der Plas et al., 2020). It is well acknowledged that intraspecific trait shift is prevalent and play a significant role in shaping plant community response to biotic and abiotic environmental perturbations (Chen et al., 2021; de Bello et al., 2021; Funk et al., 2017; Mitchell & Bakker, 2014; Siefert et al., 2015). Using traits extracted from global databases, we found that species-level leaf N, and to a lesser extent SLA, were more tightly linked to facets of stability than the LES that included LDMC but was based on species- and genus-level trait values. Thus, using species-level traits may be essential to link to ecosystem stability. However, major pathways in which nutrient addition impacted community-level traits and stability differed from those based on traits measured in the field (e.g. leaf N). While traits extracted from global databases allow for broader comparisons and generalisations across different ecosystems and regions, they may not accurately represent plant traits measured on-site. Our results suggest that caution should be taken in interpreting results based on traits extracted from global databases for ecosystem functions and stability because they may fail to capture important ecological processes induced by intraspecific trait shifts.

### Stability facets and the influencing pathways

Our results showed that nutrient addition could impact different facets of stability through different pathways. This is consistent with a previous study showing that these facets of stability are overall weakly correlated under control and nutrient addition treatments (Chen et al., 2023). Nutrient addition may impact resistance more than recovery and temporal invariability through diversity-mediated indirect effects. Past studies often investigate temporal invariability through mean and standard deviation (Hautier et al., 2015; Suonan et al., 2023). It is also well-established that global changes such as nutrient enrichment can impact community stability through changing species asynchrony (i.e. increase in biomass of some species being compensated by decrease in others) and/or population stability (Loreau & de Mazancourt, 2013). Here we disentangle the drivers underlying long-term temporal stability (i.e. temporal invariability) into two processes, resistance during and recovery after short-term extreme perturbations. In doing so, we linked plant communities’ short-term responses to long-term ones, which were often studied separately in previous studies (Donohue et al., 2013; Kéfi et al., 2019). Our results suggest that management strategies aiming at increasing resistance may be more important than those increasing recovery for maintaining temporal invariability of grassland aboveground biomass production.

## Conclusion

Our study filled an important knowledge gap by exploring the role of multifaceted plant diversity in predicting multidimensional ecosystem stability under nutrient addition. Our results suggested that intraspecific trait shift was a major driver for change in functional trait composition of LES under nutrient addition, which may further impact ecosystem stability. But functional diversity of LES was more important than functional trait composition in linking to ecosystem functioning and stability. Such diversity-mediated indirect effects, though weak, should be taken into account together with the direct effects of nutrient addition for more accurate predictions for ecosystem stability under global changes.

## Supporting information

Supporting Information for Change in functional trait diversity mediates the effects of nutrient addition on grassland stability

## Acknowledgments

We thank researchers from the NutNet who contributed data to our analysis, supplementary Table S9 lists these researchers. We thank the Minnesota Supercomputing Institute for hosting project data and the Institute on the Environment for hosting Network meetings. Nitrogen fertiliser was donated to NutNet by Crop Production Services, Loveland, CO. This experiment is funded by individual researchers at the site scale. We thank three anonymous reviewers and the associated editor for their constructive comments and suggestions to improve our manuscript.

## Funding

National Natural Science Foundation of China, grant 32122053 (SW).

National Key Research and Development Program of China, grant 2022YFF0802103 (SW).

National Science Foundation grant NSF-DEB-1042132 (ETB, EWS; for NutNet coordination and data management)

National Science Foundation grant NSF-DEB-1234162 (ETB, EWS; for Long-Term Ecological Research at Cedar Creek).

National Science Foundation grant NSF-DEB-1831944 (ETB, EWS; for Long-Term Ecological Research at Cedar Creek)

## Competing interests

Authors declare that they have no competing interests. Yann Hautier is an Associate Editor of Journal of Ecology, but took no part in the peer review and decision-making processes for this paper.

## Author contributions (see Table S10 for more details for contribution of each author)

Qingqing Chen, Shaopeng Wang, and Yann Hautier conceived the ideas; Qingqing Chen, Shaopeng Wang, Forest Isbell, Siddharth Bharath, Juan Alberti, and Yann Hautier designed methodology; Eric W. Seabloom, Elizabeth T. Borer, Jonathan D. Bakker, Christiane Roscher, Sally A. Power, Carly Stevens, Anne Ebeling, Carla Nogueira, Maria C Caldeira, Andrew S. MacDougall, Joslin L. Moore, Sumanta Bagchi, Michelle Tedder, Kevin Kirkman, collected the data; Qingqing Chen analysed the data; Qingqing Chen led the writing of the manuscript. All authors contributed critically to the drafts and gave final approval for publication.

## Data and materials availability

Data are stored in Figshare (https://doi.org/10.6084/m9.figshare.26489773.v1). The R codes for compiling traits from global databases are deposited in the Github (https://github.com/chqq365/plant-traits-compiled-for-NutNet), and archived through Zenodo (https://doi.org/10.5281/zenodo.13204824). The R codes used to produce results are also deposited in the Github (https://github.com/chqq365/Plant-traits-and-multidimensional-stability), and archived through Zenodo (https://doi.org/10.5281/zenodo.13206421).

## Notes

### Competing Interest Statement

The authors have declared no competing interest.

### Summary of Updates

Add missing citations; add links for data and R codes associated with the manuscript. Improve visualization of the graph.

## References

Bazzichetto, M., Sperandii, M. G., Penone, C., Keil, P., Allan, E., Lepš, J., Prati, D., Fischer, M., Bolliger, R., Gossner, M. M., & de Bello, F. (2024). Biodiversity promotes resistance but dominant species shape recovery of grasslands under extreme drought. Journal of Ecology, n/a(n/a). 10.1111/1365-2745.14288

Borer, E. T., Grace, J. B., Harpole, W. S., MacDougall, A. S., & Seabloom, E. W. (2017). A decade of insights into grassland ecosystem responses to global environmental change. Nature Ecology and Evolution, 1(5), 1–7. 10.1038/s41559-017-0118

Borer, E. T., Harpole, W. S., Adler, P. B., Lind, E. M., Orrock, J. L., Seabloom, E. W., & Smith, M. D. (2014). Finding generality in ecology: A model for globally distributed experiments. Methods in Ecology and Evolution, 5(1), 65–73. 10.1111/2041-210X.12125

Chase, J. M., McGill, B. J., McGlinn, D. J., May, F., Blowes, S. A., Xiao, X., Knight, T. M., Purschke, O., & Gotelli, N. J. (2018). Embracing scale-dependence to achieve a deeper understanding of biodiversity and its change across communities. Ecology Letters, 21(11), 1737–1751. 10.1111/ele.13151

Chen, Q., Smit, C., Pen, I., & Olff, H. (2021). Small herbivores and abiotic heterogeneity promote trait variation of a saltmarsh plant in local communities. PeerJ, 9, e12633. 10.7717/PEERJ.12633

Chen, Q., Wang, S., Borer, E. T., Bakker, J. D., Seabloom, E. W., Harpole, W. S., Eisenhauer, N., Lekberg, Y., Buckley, Y. M., Catford, J. A., Roscher, C., Donohue, I., Power, S. A., Daleo, P., Ebeling, A., Knops, J. M. H., Martina, J. P., Eskelinen, A., Morgan, J. W., … Hautier, Y. (2023). Multidimensional responses of grassland stability to eutrophication. Nature Communications, 14(1), Article 1. 10.1038/s41467-023-42081-0

Chen, Q., Wang, S., Seabloom, E. W., MacDougall, A. S., Borer, E. T., Bakker, J. D., Donohue, I., Knops, J. M. H., Morgan, J. W., Carroll, O., Crawley, M., Bugalho, M. N., Power, S. A., Eskelinen, A., Virtanen, R., Risch, A. C., Schütz, M., Stevens, C., Caldeira, M. C., … Hautier, Y. (2022). Nutrients and herbivores impact grassland stability across spatial scales through different pathways. Global Change Biology, 28(8), 2678–2688. 10.1111/gcb.16086

Chen, Q., Wang, S., Seabloom, E.W., Forest, I., Borer, E.T., Bakker, J.D., et al. (2024). Data for Change in functional trait diversity mediates the effects of nutrient addition on grassland stability. Figshare. Dataset. 10.6084/m9.figshare.26489773.v1.

Chen, Q. (2024). chqq365/Plant-traits-and-multidimensional-stability: R codes to support Change in functional trait diversity mediates the effects of nutrient addition on grassland stability; 10.5281/zenodo.13204824.

Chen, Q. (2024). chqq365/plant-traits-compiled-for-NutNet: Functional traits compilied from global databases for NutNet species; 10.5281/zenodo.13206421.

Craven, D., Polley, H. W., & Wilsey, B. (2018). Multiple facets of biodiversity drive the diversity – stability relationship. Nature Ecology & Evolution, 2, 1579–1587. 10.1038/s41559-018-0647-7.Rights

de Bello, F., Lavorel, S., Hallett, L. M., Valencia, E., Garnier, E., Roscher, C., Conti, L., Galland, T., Goberna, M., Májeková, M., Montesinos-Navarro, A., Pausas, J. G., Verdú, M., E-Vojtkó, A., Götzenberger, L., & Lepš, J. (2021). Functional trait effects on ecosystem stability: Assembling the jigsaw puzzle. Trends in Ecology and Evolution, 1–15. 10.1016/j.tree.2021.05.001

Donohue, I., Petchey, O. L., Montoya, J. M., Jackson, A. L., Mcnally, L., Viana, M., Healy, K., Lurgi, M., O’Connor, N. E., & Emmerson, M. C. (2013). On the dimensionality of ecological stability. Ecology Letters, 16(4), 421–429. 10.1111/ele.12086

Duodu, G. O., Goonetilleke, A., Allen, C., & Ayoko, G. (2015). Determination of refractive and volatile elements in sediment using laser ablation inductively coupled plasma mass spectrometry. Anal. Chim. Acta, 898, 19–27.

Falster, D., Gallagher, R., Wenk, E. H., Wright, I. J., Indiarto, D., Andrew, S. C., Baxter, C., Lawson, J., Allen, S., Fuchs, A., Monro, A., Kar, F., Adams, M. A., Ahrens, C. W., Alfonzetti, M., Angevin, T., Apgaua, D. M. G., Arndt, S., Atkin, O. K., … Ziemińska, K. (2021). AusTraits, a curated plant trait database for the Australian flora. Scientific Data, 8(1), 1–20. 10.1038/s41597-021-01006-6

Firn, J., McGree, J. M., Harvey, E., Flores-Moreno, H., Schütz, M., Buckley, Y. M., Borer, E. T., Seabloom, E. W., La Pierre, K. J., MacDougall, A. M., Prober, S. M., Stevens, C. J., Sullivan, L. L., Porter, E., Ladouceur, E., Allen, C., Moromizato, K. H., Morgan, J. W., Harpole, W. S., … Risch, A. C. (2019). Leaf nutrients, not specific leaf area, are consistent indicators of elevated nutrient inputs. Nature Ecology and Evolution, 3(3), 400–406. 10.1038/s41559-018-0790-1

Funk, J. L., Larson, J. E., Ames, G. M., Butterfield, B. J., Cavender-Bares, J., Firn, J., Laughlin, D. C., Sutton-Grier, A. E., Williams, L., & Wright, J. (2017). Revisiting the Holy Grail: Using plant functional traits to understand ecological processes. Biological Reviews, 92(2), 1156– 1173. 10.1111/brv.12275

Galloway, J. N., Bleeker, A., & Erisman, J. W. (2021). The Human Creation and Use of Reactive Nitrogen: A Global and Regional Perspective. Annual Review of Environment and Resources, 46(1), 255–288. 10.1146/annurev-environ-012420-045120

Garnier, Cortez, J., Billès, G., Navas, M.-L., Roumet, C., Debussche, M., Laurent, G., Blanchard, A., Aubry, D., Bellmann, A., Neill, C., & Toussaint, J.-P. (2004). Plant Functional Markers Capture Ecosystem Properties During Secondary Succession. Ecology, 85(9), 2630–2637. 10.1890/03-0799

Harpole, W. S., Potts, D. L., & Suding, K. N. (2007). Ecosystem responses to water and nitrogen amendment in a California grassland. Global Change Biology, 13(11), 2341–2348. 10.1111/j.1365-2486.2007.01447.x

Harris, I., Osborn, T. J., Jones, P., & Lister, D. (2020). Version 4 of the CRU TS monthly high-resolution gridded multivariate climate dataset. Scientific Data, 7(1), 1–18. 10.1038/s41597-020-0453-3

Hautier, Y., Tilman, D., Isbell, F., Seabloom, E. W., Borer, E. T., & Reich, P. B. (2015). Anthropogenic environmental changes affect ecosystem stability via biodiversity. Science, 348(6232), 336–340. 10.1126/science.aaa1788

IPCC. (2023): Summary for Policymakers. In: Climate Change 2023: Synthesis Report. Contribution of Working Groups I, II and III to the Sixth Assessment Report of the Intergovernmental Panel on Climate Change [Core Writing Team, H. Lee and J. Romero (eds.)]. IPCC, Geneva, Switzerland, pp. 1–34, doi: 10.59327/IPCC/AR6-9789291691647.001. 10.1016/j.tree.2023.03.001

Isbell, F., Craven, D., Connolly, J., Loreau, M., Schmid, B., Beierkuhnlein, C., Bezemer, T. M., Bonin, C., Bruelheide, H., de Luca, E., Ebeling, A., Griffin, J. N., Guo, Q., Hautier, Y., Hector, A., Jentsch, A., Kreyling, J., Lanta, V., Manning, P., … Eisenhauer, N. (2015). Biodiversity increases the resistance of ecosystem productivity to climate extremes. Nature, 526(7574), 574–577. 10.1038/nature15374

Ives, A. R., & Carpenter, S. R. (2007). Supporting file for Stability and diversity of ecosystems. Science, 317(5834), 58–62. 10.1126/science.1133258

Jung, V., Albert, C. H., Violle, C., Kunstler, G., Loucougaray, G., & Spiegelberger, T. (2014). Intraspecific trait variability mediates the response of subalpine grassland communities to extreme drought events. Journal of Ecology, 102(1), 45–53. 10.1111/1365-2745.12177

Kaspari, M. (2021). The Invisible Hand of the Periodic Table: How Micronutrients Shape Ecology. Annual Review of Ecology, Evolution, and Systematics, 52(1), 199–219. 10.1146/annurev-ecolsys-012021-090118

Kattge, J., Bönisch, G., Díaz, S., Lavorel, S., Prentice, I. C., Leadley, P., Tautenhahn, S., Werner, G. D. A., Aakala, T., Abedi, M., Acosta, A. T. R., Adamidis, G. C., Adamson, K., Aiba, M., Albert, C. H., Alcántara, J. M., Alcázar C, C., Aleixo, I., Ali, H., … Wirth, C. (2020). TRY plant trait database – enhanced coverage and open access. Global Change Biology, 26(1), 119–188. 10.1111/gcb.14904

Kéfi, S., Domínguez-García, V., Donohue, I., Fontaine, C., Thébault, E., & Dakos, V. (2019). Advancing our understanding of ecological stability. Ecology Letters, 22(9), 1349–1356. 10.1111/ele.13340

Laliberte, E., & Legendre, P. (2010). A distance-based framework for measuring functional diversity from multiple traits. Ecology, 91(1), 299–305. 10.1890/08-2244.1

Laliberté, E., Legendre, P., & Shipley, B. (2014). Package ‘FD.’ 85–91.

Lê, S., Josse, J., & Husson, F. (2008). FactoMineR: An R package for multivariate analysis. Journal of Statistical Software, 25(1), 1–18. 10.18637/jss.v025.i01

Lefcheck, J. S. (2016). piecewiseSEM: Piecewise structural equation modelling in r for ecology, evolution, and systematics. Methods in Ecology and Evolution, 7(5), 573–579. 10.1111/2041-210X.12512

Lepš, J., de Bello, F., Šmilauer, P., & Doležal, J. (2011). Community trait response to environment: Disentangling species turnover vs intraspecific trait variability effects. Ecography, 34(5), 856–863.

Loreau, M., & de Mazancourt, C. (2013). Biodiversity and ecosystem stability: A synthesis of underlying mechanisms. Ecology Letters, 16(SUPPL.1), 106–115. 10.1111/ele.12073

Maitner, B. S., Boyle, B., Casler, N., Condit, R., Donoghue II, J., Durán, S. M., Guaderrama, D., Hinchliff, C. E., Jørgensen, P. M., Kraft, N. J. B., McGill, B., Merow, C., Morueta-Holme, N., Peet, R. K., Sandel, B., Schildhauer, M., Smith, S. A., Svenning, J.-C., Thiers, B., … Enquist, B. J. (2018). The bien r package: A tool to access the Botanical Information and Ecology Network (BIEN) database. Methods in Ecology and Evolution, 9(2), 373–379. 10.1111/2041-210X.12861

Mariotte, P., Vandenberghe, C., Kardol, P., Hagedorn, F., & Buttler, A. (2013). Subordinate plant species enhance community resistance against drought in semi-natural grasslands. Journal of Ecology, 101(3), 763–773. 10.1111/1365-2745.12064

Meng, B., Li, J., Maurer, G. E., Zhong, S., Yao, Y., Yang, X., Collins, S. L., & Sun, W. (2021). Nitrogen addition amplifies the nonlinear drought response of grassland productivity to extended growing-season droughts. Ecology, 102(11), e03483. 10.1002/ecy.3483

Mitchell, R. M., & Bakker, J. D. (2014). Intraspecific Trait Variation Driven by Plasticity and Ontogeny in Hypochaeris radicata. PLOS ONE, 9(10), e109870. 10.1371/journal.pone.0109870

Oram, N. J., De Deyn, G. B., Bodelier, P. L. E., Cornelissen, J. H. C., van Groenigen, J. W., & Abalos, D. (2020). Plant community flood resilience in intensively managed grasslands and the role of the plant economic spectrum. Journal of Applied Ecology, 57(8), 1524–1534. 10.1111/1365-2664.13667

Pérez-Harguindeguy, N., Díaz, S., Garnier, E., Lavorel, S., Poorter, H., Jaureguiberry, P., Bret-Harte, M. S., Cornwell, W. K., Craine, J. M., Gurvich, D. E., Urcelay, C., Veneklaas, E. J., Reich, P. B., Poorter, L., Wright, I. J., Ray, P., Enrico, L., Pausas, J. G., de Vos, A. C., … Cornelissen, J. H. C. (2016). Corrigendum to: New handbook for standardised measurement of plant functional traits worldwide. Australian Journal of Botany, 64(8), 715. 10.1071/bt12225_co

Pichon, N. A., Cappelli, S. L., & Allan, E. (2022). Intraspecific trait changes have large impacts on community functional composition but do not affect ecosystem function. Journal of Ecology. 10.1111/1365-2745.13827

Pimm, S. L. (1984). The complexity and stability of ecosystems. Nature, 307(5949), 321–326. 10.1038/307321a0

Pinheiro, J., Bates, D., DebRoy, S., Sarkar, D., & Team, R.-C. (2017). nlme: Linear and Nonlinear Mixed Effects Models. R Package Version 3.1-131. https://cran.r-project.org/package=nlme

R Core Team. (2022). R: A language and environment for statistical computing. [Computer software]. R Foundation for Statistical Computing.

Reich, P. B. (2014). The world-wide “fast-slow” plant economics spectrum: A traits manifesto. Journal of Ecology, 102(2), 275–301. 10.1111/1365-2745.12211

Robinson, T. M. P., La Pierre, K. J., Vadeboncoeur, M. A., Byrne, K. M., Thomey, M. L., & Colby, S. E. (2013). Seasonal, not annual precipitation drives community productivity across ecosystems. Oikos, 122(5), 727–738. 10.1111/j.1600-0706.2012.20655.x

Sala, O. E., Gherardi, L. A., Reichmann, L., Jobbágy, E., & Peters, D. (2012). Legacies of precipitation fluctuations on primary production: Theory and data synthesis. Philosophical Transactions of the Royal Society B: Biological Sciences, 367(1606), 3135–3144. 10.1098/rstb.2011.0347

Sardans, J., & Peñuelas, J. (2015). Potassium: A neglected nutrient in global change. Global Ecology and Biogeography, 24(3), 261–275. 10.1111/geb.12259

Schnabel, F., Liu, X., Kunz, M., Barry, K. E., Bongers, F. J., Bruelheide, H., Fichtner, A., Härdtle, W., Li, S., Pfaff, C. T., Schmid, B., Schwarz, J. A., Tang, Z., Yang, B., Bauhus, J., Von Oheimb, G., Ma, K., & Wirth, C. (2021). Species richness stabilizes productivity via asynchrony and drought-tolerance diversity in a large-scale tree biodiversity experiment. Science Advances, 7(51), 11–13. 10.1126/sciadv.abk1643

Siefert, A., & Ritchie, M. E. (2016). Intraspecific trait variation drives functional responses of old-field plant communities to nutrient enrichment. Oecologia, 181(1), 245–255. 10.1007/s00442-016-3563-z

Siefert, A., Violle, C., Chalmandrier, L., Albert, C. H., Taudiere, A., Fajardo, A., Aarssen, L. W., Baraloto, C., Carlucci, M. B., Cianciaruso, M. V., de L. Dantas, V., de Bello, F., Duarte, L. D. S., Fonseca, C. R., Freschet, G. T., Gaucherand, S., Gross, N., Hikosaka, K., Jackson, B., … Wardle, D. A. (2015). A global meta-analysis of the relative extent of intraspecific trait variation in plant communities. Ecology Letters, 18(12), 1406–1419. 10.1111/ele.12508

Southon, G. E., Green, E. R., Jones, A. G., Barker, C. G., & Power, S. A. (2012). Long-term nitrogen additions increase likelihood of climate stress and affect recovery from wildfire in a lowland heath. Global Change Biology, 18(9), 2824–2837. 10.1111/j.1365-2486.2012.02732.x

Sun, J., Liu, W., Pan, Q., Zhang, B., Lv, Y., Huang, J., & Han, X. (2022). Positive legacies of severe droughts in the Inner Mongolia grassland. Science Advances, 8(47), eadd6249. 10.1126/sciadv.add6249

Suonan, J., Lu, X., Li, X., Hautier, Y., & Wang, C. (2023). Nitrogen addition strengthens the stabilizing effect of biodiversity on productivity by increasing plant trait diversity and species asynchrony in the artificial grassland communities. Frontiers in Plant Science, 14. 10.3389/fpls.2023.1301461

Tatarko, A. R., & Knops, J. M. H. (2018). Nitrogen addition and ecosystem functioning: Both species abundances and traits alter community structure and function. Ecosphere, 9(1). 10.1002/ecs2.2087

Tilman, D. (1996). Biodiversity: Population Versus ecosystem stability. Ecology, 77(2), 350–363.

Tilman, D., Reich, P. B., & Knops, J. M. H. (2006). Biodiversity and ecosystem stability in a decade-long grassland experiment. Nature, 441(7093), Article 7093. 10.1038/nature04742

van der Plas, F., Schröder-Georgi, T., Weigelt, A., Barry, K., Meyer, S., Alzate, A., Barnard, R. L., Buchmann, N., de Kroon, H., Ebeling, A., Eisenhauer, N., Engels, C., Fischer, M., Gleixner, G., Hildebrandt, A., Koller-France, E., Leimer, S., Milcu, A., Mommer, L., … Wirth, C. (2020). Plant traits alone are poor predictors of ecosystem properties and long-term ecosystem functioning. Nature Ecology & Evolution, 4(12), 1602–1611. 10.1038/s41559-020-01316-9

Wheeler, Brassil, C. E., & Knops, J. M. H. (2022). Functional traits’ annual variation exceeds nitrogen-driven variation in grassland plant species. Ecology, 104(2), e3886. 10.1002/ecy.3886

Wheeler, M. M., Collins, S. L., Grimm, N. B., Cook, E. M., Clark, C., Sponseller, R. A., & Hall, S. J. (2021). Water and nitrogen shape winter annual plant diversity and community composition in near-urban Sonoran Desert preserves. Ecological Monographs, 91(3). 10.1002/ecm.1450

Wright, I. J., Reich, P. B., Westoby, M., Ackerly, D. D., Baruch, Z., Bongers, F., Cavender-Bares, J., Chapin, T., Cornellssen, J. H. C., Diemer, M., Flexas, J., Garnier, E., Groom, P. K., Gulias, J., Hikosaka, K., Lamont, B. B., Lee, T., Lee, W., Lusk, C., … Villar, R. (2004). The worldwide leaf economics spectrum. Nature, 428(6985), 821–827. 10.1038/nature02403

Xu, Q., Yang, X., Song, J., Ru, J., Xia, J., Wang, S., Wan, S., & Jiang, L. (2022). Nitrogen enrichment alters multiple dimensions of grassland functional stability via changing compositional stability. Ecology Letters, 25(12), 2713–2725. 10.1111/ele.14119

Yuan, Z., Jiang, S., Sheng, H., Liu, X., Hua, H., Liu, X., & Zhang, Y. (2018). Human Perturbation of the Global Phosphorus Cycle: Changes and Consequences. Environmental Science & Technology, 52(5), 2438–2450. 10.1021/acs.est.7b03910

Zhou, X., Guo, Z., Zhang, P., & Du, G. (2018). Shift in community functional composition following nitrogen fertilization in an alpine meadow through intraspecific trait variation and community composition change. Plant and Soil, 431(1–2), 289–302. 10.1007/S11104-018-3771-X/TABLES/5

