## Supporting Information for Change in functional trait diversity mediates the effects of nutrient addition on grassland stability for "Change in functional trait diversity mediates the effects of nutrient addition on grassland stability"

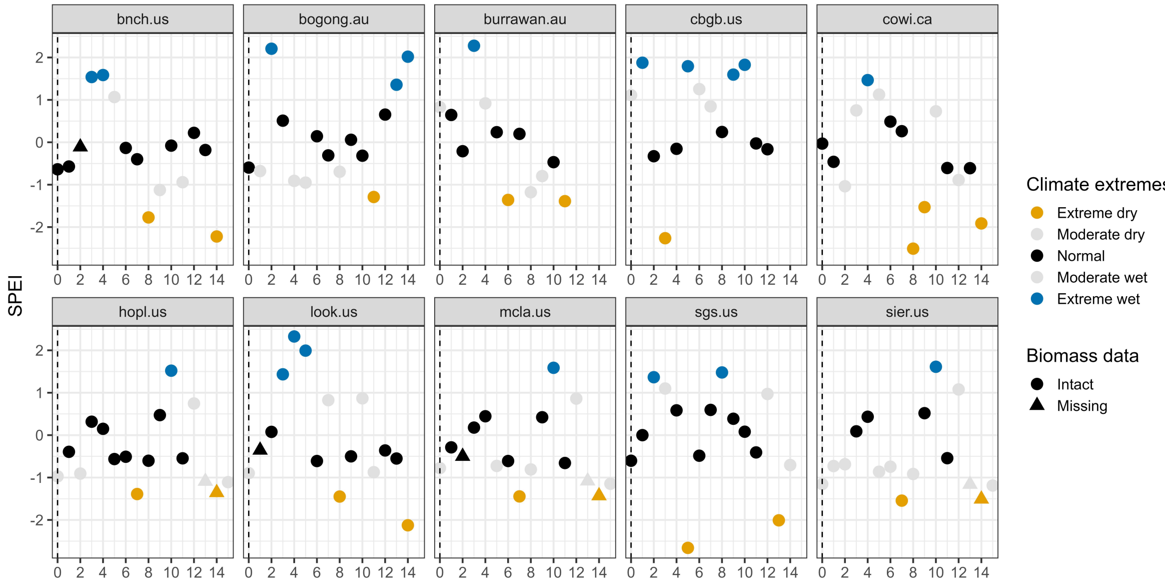


**Fig. S1 Categorize dry and wet growing seasons at each site during experimental years at 10 NutNet sites.** When including moderate extreme growing seasons, for some years at some sites, dry and wet growing seasons occurred consecutively, limiting our ability to disentangle resistance and recovery. Therefore, we focus on extreme dry and wet growing seasons to reduce confounding effects. Also, more extreme climate conditions may allow us to detect greater changes in aboveground biomass. Resistance and recovery present below were based on extreme growing seasons.

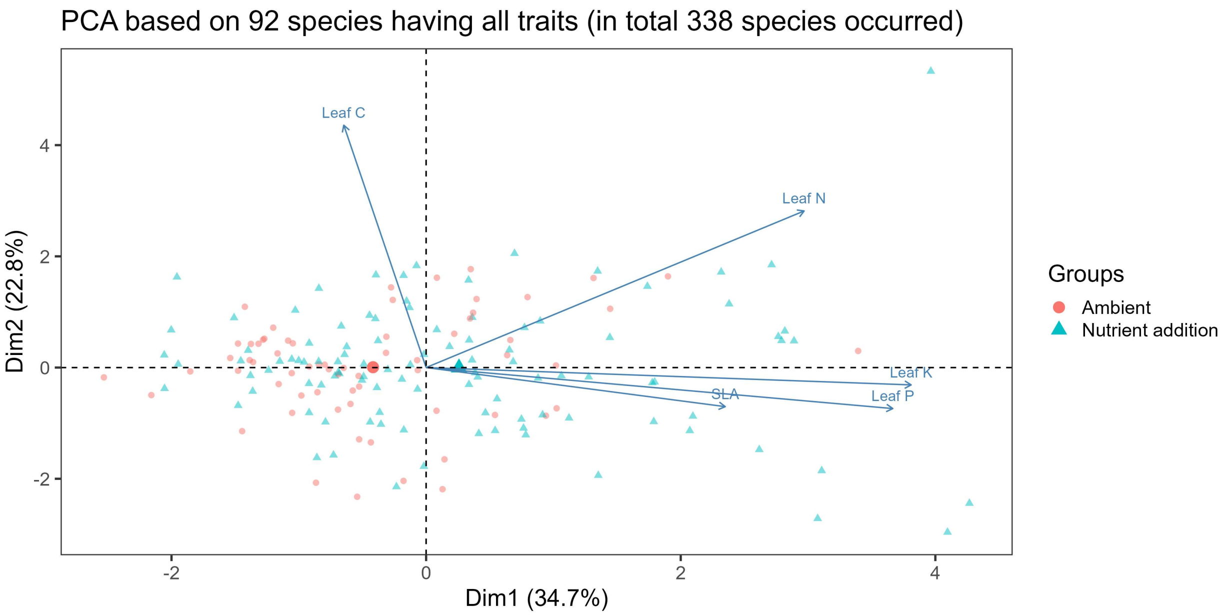


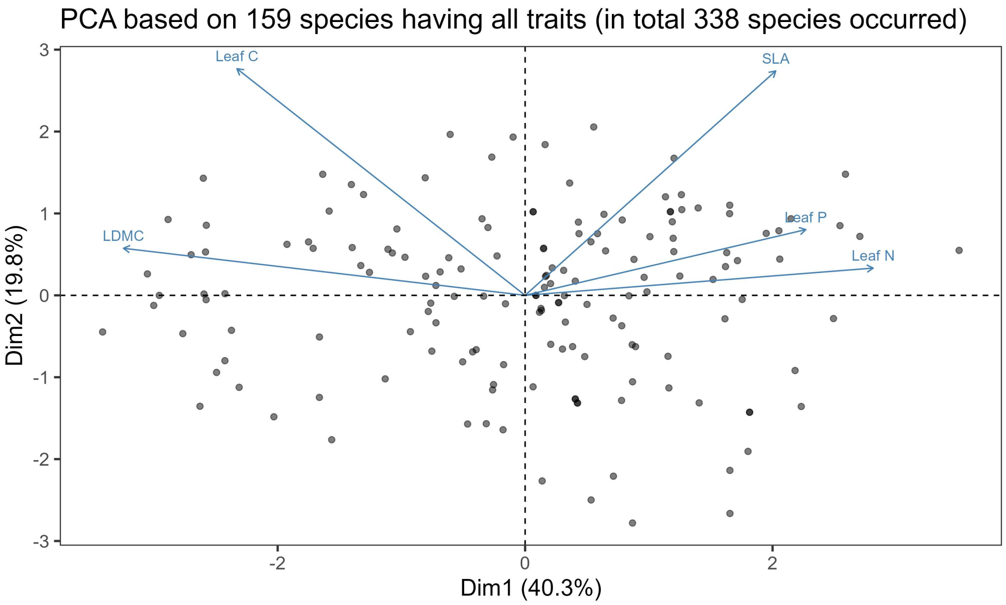


**Fig. S2 PCA using five leaf traits for species from the ambient and nutrient addition conditions across 10 sites (upper panel) and based on trait extracted from global trait databases (lower panel).**

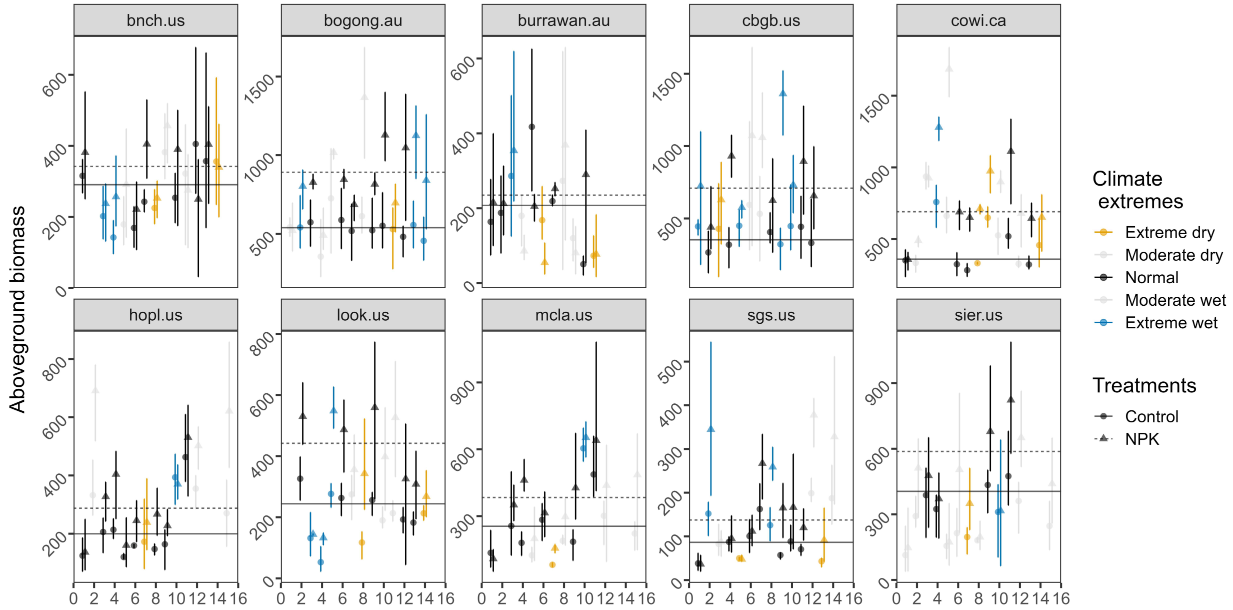


**Fig. S3 Aboveground biomass (g m$^{-2}$) over the experimental years in control and nutrient addition treatments at each site.** Climate extremes correspond to Fig. S1. Dots indicate raw live biomass data averaged over three blocks in each year at each site. Lines indicate average live biomass over normal growing seasons over three blocks at each site. Error bars are 95% bootstrapped confidence interval.

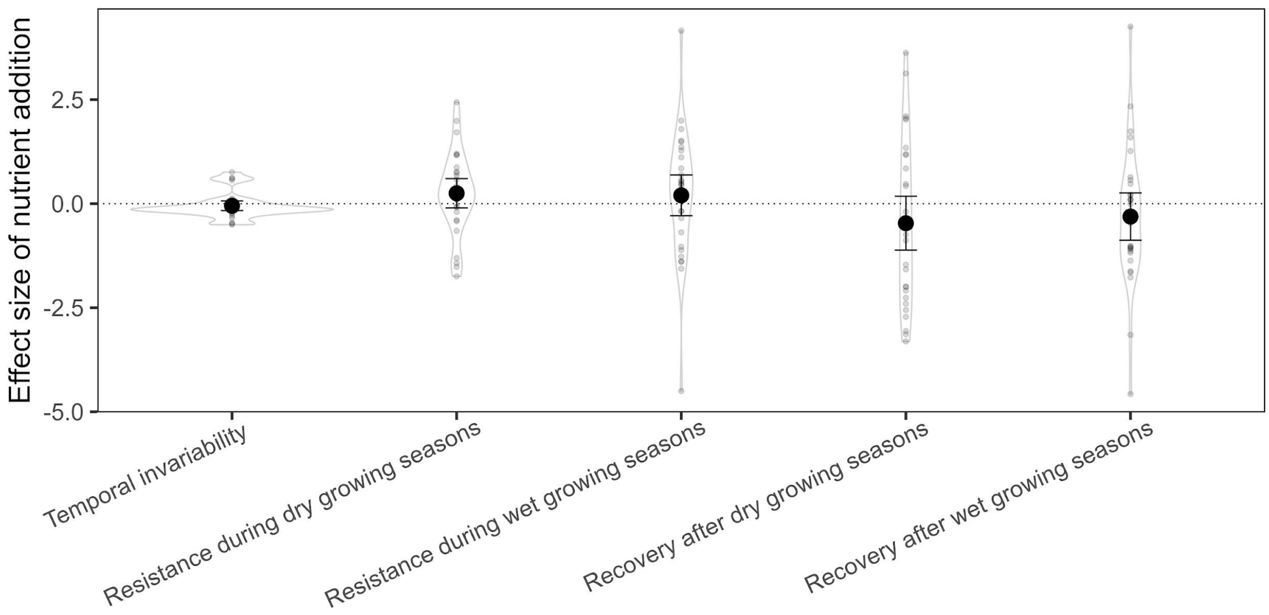


**Fig. S4. Effects of nutrient addition on facets of stability.** Resistance and recovery are quantified using normal levels and deviation from normal levels shown in Fig. 3. Small points are effects of nutrient addition on each stability facet from each block at each site. Large black points are mean over experimental years and over 10 sites estimated from linear mixed effect models, error bars are 95% confidence intervals. Violin shapes show distribution of values. All stability facets are on a log scale. See Table S6 for test statistics.

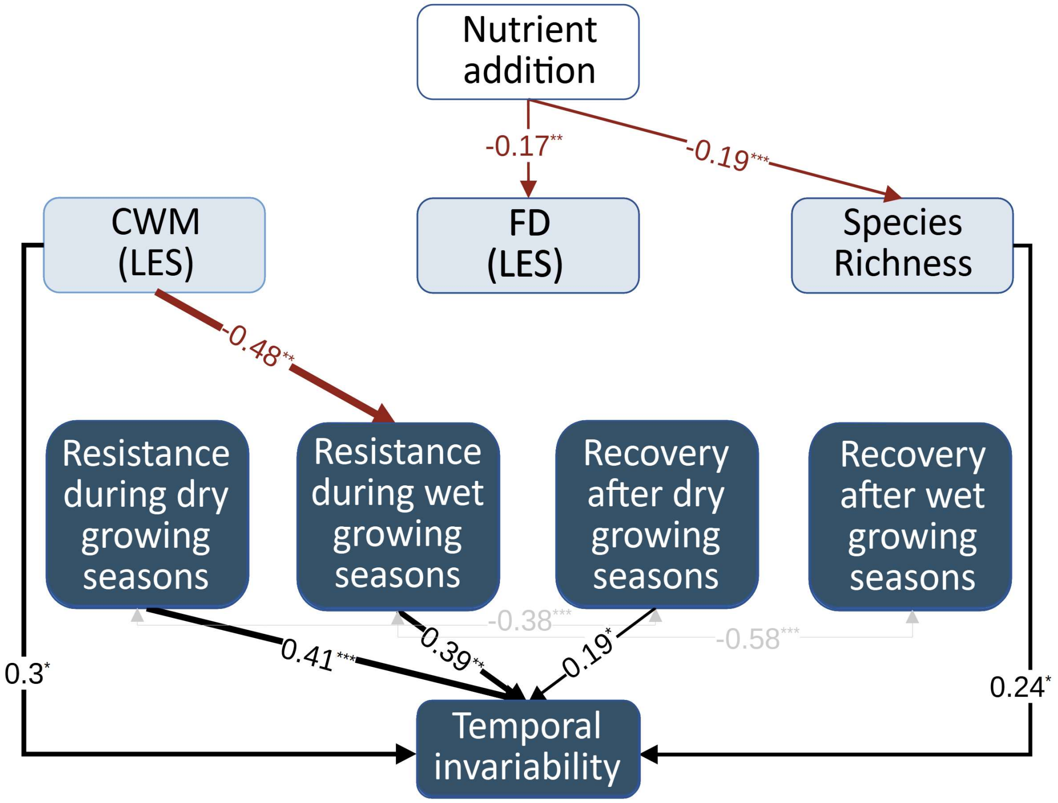


**Fig. S5 Direct and diversity-mediated effects of nutrient addition on facets of stability.** CWM and FD were calculated based on constructed leaf economic spectrum (LES) using species- and genus-level filled traits from global trait databases. White box represents nutrient addition by NPK, blue boxes represent measured variables, and arrows represent relationships among variables. The displayed numbers are standardized path coefficients. The width of the arrows indicates the strength of the pathways. Line color represents positive (black) and negative (red) effects. Non-significant paths are not shown. Grey lines and text show correlated errors. Asterisks indicate significant paths: * $p \leq 0.1$; ** $p \leq 0.05$; *** $p \leq 0.001$. All stability facets were on the log scale to improve normality and homogeneity of variance. See Table S7 for variance explained (R2) for each component model and overall goodness of model fit. See Table S8 for a summary of the direct and diversity-mediated indirect effects of nutrient addition on facets of stability.

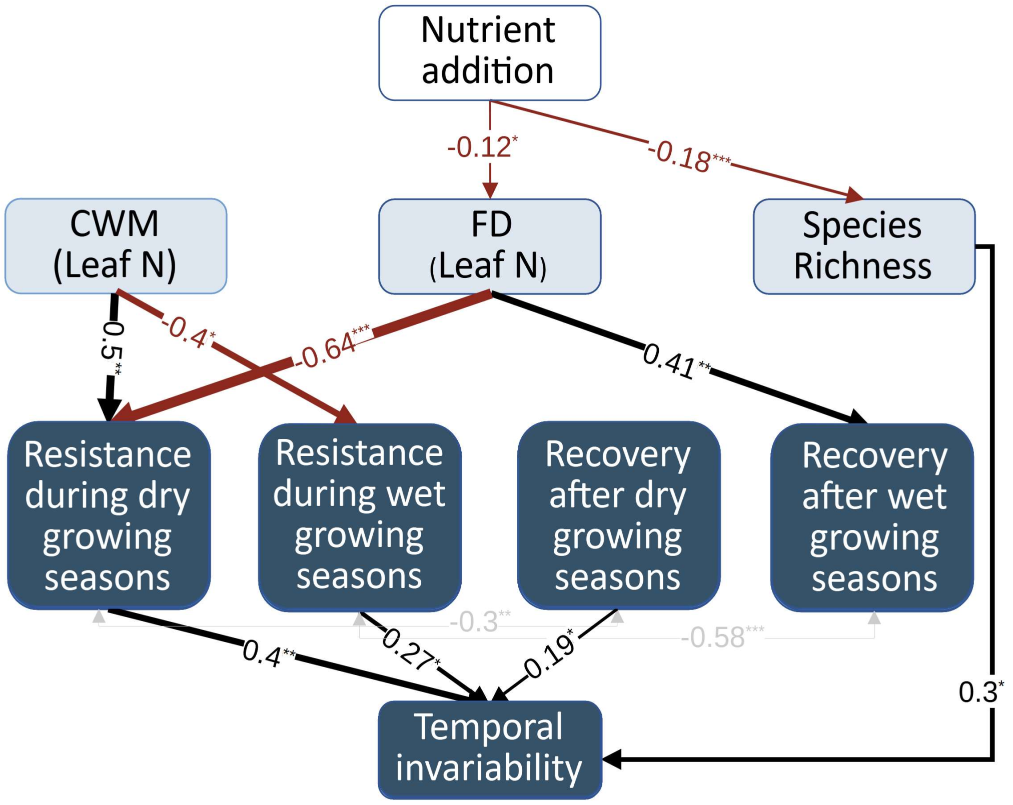


**Fig. S6 Direct and diversity-mediated effects of nutrient addition on facets of stability.** CWM and FD were calculated based on species-level leaf N from global trait databases. White box represents nutrient addition by NPK, blue boxes represent measured variables, and arrows represent relationships among variables. The displayed numbers are standardized path coefficients. The width of the arrows indicates the strength of the pathways. Line color represents positive (black) and negative (red) effects. Non-significant paths are not shown. Grey lines and text show correlated errors. Asterisks indicate significant paths: * $p \leq 0.1$; ** $p \leq 0.05$; *** $p \leq 0.001$. All stability facets were on the log scale to improve normality and homogeneity of variance. See Table S7 for variance explained (R2) for each component model and overall goodness of model fit. See Table S8 for a summary of the direct and diversity-mediated indirect effects of nutrient addition on facets of stability.

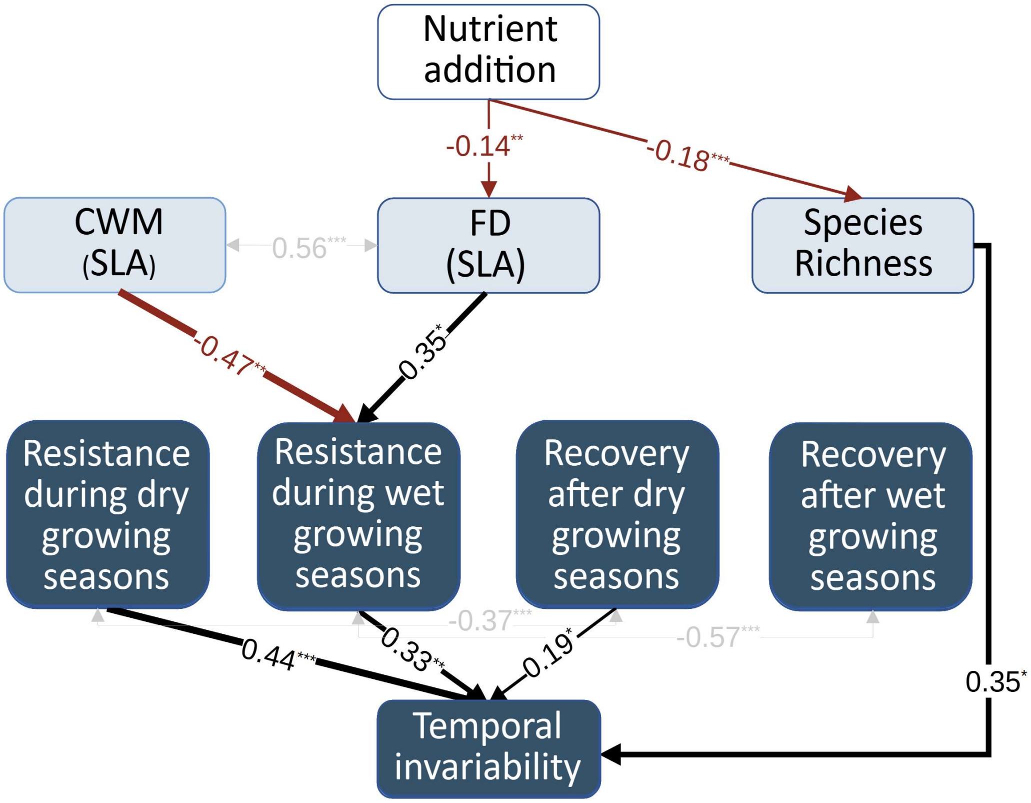


**Fig. S7 Direct and diversity-mediated effects of nutrient addition on facets of stability.** CWM and FD were calculated based on species-level SLA from global trait databases. White box represents nutrient addition by NPK, blue boxes represent measured variables, and arrows represent relationships among variables. The displayed numbers are standardized path coefficients. The width of the arrows indicates the strength of the pathways. Line color represents positive (black) and negative (red) effects. Non-significant paths are not shown. Grey lines and text show correlated errors. Asterisks indicate significant paths: * $p \leq 0.1$; ** $p \leq 0.05$; *** $p \leq 0.001$. All stability facets were on the log scale to improve normality and homogeneity of variance. See Table S7 for variance explained (R2) for each component model and overall goodness of model fit. See Table S8 for a summary of the direct and diversity-mediated indirect effects of nutrient addition on facets of stability.

**Table S1 sites selected and their geolocation and experimental years used.** Growing season refers to from the start to the end of growing month at each site. Growing season is recorded by site PIs. Water balance is calculated as the mean of precipitation - potential evaporation over growing seasons from 2005 to 2022 at each site, positive values suggest wet sites, negative values suggest dry sites.

| Site_code | Habitat | Continent | Latitude | Longitude | Growing Season | Water Balance | First Experimental Year | Year used |
|---|---|---|---|---|---|---|---|---|
| bnch.us | montane grassland | North America | 44.28 | -121.97 | 4-8 | -0.10 | 2008 | 1,3-11 |
| bogong.au | alpine grassland | Australia | -36.87 | 147.25 | 10-1 | -0.16 | 2010 | 1-14 |
| burrawan.au | semiarid grassland | Australia | -27.74 | 151.14 | 10-5 | -0.04 | 2009 | 1-11 |
| cbgb.us | tallgrass prairie | North America | 41.79 | -93.39 | 5-10 | 0.40 | 2010 | 1-12 |
| cowi.ca | old field | North America | 48.81 | -123.63 | 4-7 | -0.21 | 2008 | 1-14 |
| hopl.us | annual grassland | North America | 39.01 | -123.06 | 11-4 | -0.20 | 2008 | 1-15 |
| look.us | montane grassland | North America | 44.21 | -122.13 | 3-8 | 0.07 | 2008 | 2-14 |
| mcla.us | annual grassland | North America | 38.86 | -122.41 | 11-4 | -0.22 | 2008 | 1,3-12 |
| sgs.us | shortgrass prairie | North America | 40.82 | -104.77 | 4-8 | -0.20 | 2008 | 1-14 |
| sier.us | annual grassland | North America | 39.24 | -121.28 | 11-4 | -0.29 | 2008 | 1-15 |

**Table S2 Information for leaf traits data at 10 sites used.** Community-weighted mean (CWM) of leaf economic spectrum (LES) are present here.

| Site_code | Years for trait measurement | Proportional cover of species with traits under Control and NPK | Proportional cover of shared species with traits under Control and NPK | Shift in CWM based on all species | Shift in CWM based on shared species | Shift in CWM due to replacement among existing species | Shift in CWM due to intraspecific trait shift |
|---|---|---|---|---|---|---|---|
| bnch.us | 4 | 0.52/0.67 | 0.51/0.55 | 0.52 | 0.46 | -0.04 | 0.50 |
| bogong.au | 2 | 0.87/0.84 | 0.79/0.67 | 0.73 | 0.55 | 0.22 | 0.33 |
| burrawan.au | 3 | 0.88/0.84 | 0.85/0.78 | 0.65 | 0.67 | 0.14 | 0.53 |
| cbgb.us | 3 | 0.8/0.89 | 0.56/0.65 | 0.37 | 0.44 | 0.16 | 0.28 |
| cowi.ca | 4 | 0.92/0.99 | 0.9/0.99 | 0.24 | 0.25 | -0.04 | 0.30 |
| hopl.us | 4 | 0.52/0.82 | 0.4/0.54 | 0.93 | 0.45 | 0.10 | 0.35 |
| look.us | 4 | 0.64/0.72 | 0.52/0.64 | -0.19 | -0.55 | -0.56 | 0.01 |
| mcla.us | 4 | 0.82/0.87 | 0.64/0.72 | 0.38 | 0.36 | 0.08 | 0.28 |
| sgs.us | 4 | 0.65/0.7 | 0.61/0.64 | 0.79 | 0.73 | 0.08 | 0.65 |
| sier.us | 4 | 0.62/0.88 | 0.6/0.56 | 0.51 | 0.69 | -0.01 | 0.71 |

**Table S3. Proportional cover of communities with trait values extracted from global databases under control and nutrient addition by NPK per each site.**

| Trait name | site_code | Proportional cover of species with traits under Control and NPK |
|---|---|---|
| LDMC | bnch.us | 0.23/0.39 |
| LDMC | bogong.au | 0.11/0.16 |
| LDMC | burrawan.au | NA/0.06 |
| LDMC | cbgb.us | 0.93/0.87 |
| LDMC | cowi.ca | 0.97/1 |
| LDMC | hopl.us | 0.73/0.89 |

| Trait name | site_code | Proportional cover of species with traits under Control and NPK |
|---|---|---|
| LDMC | look.us | 0.36/0.5 |
| LDMC | mcla.us | 0.95/0.92 |
| LDMC | sgs.us | 0.64/0.63 |
| LDMC | sier.us | 0.91/0.96 |
| Leaf C | bnch.us | 0.25/0.44 |
| Leaf C | bogong.au | 0.61/0.6 |
| Leaf C | burrawan.au | 0.88/0.83 |
| Leaf C | cbgb.us | 0.93/0.9 |
| Leaf C | cowi.ca | 0.93/0.98 |
| Leaf C | hopl.us | 0.8/0.89 |
| Leaf C | look.us | 0.26/0.33 |
| Leaf C | mcla.us | 0.99/0.97 |
| Leaf C | sgs.us | 0.45/0.45 |
| Leaf C | sier.us | 0.89/0.95 |
| Leaf N | bnch.us | 0.31/0.38 |
| Leaf N | bogong.au | 0.62/0.61 |
| Leaf N | burrawan.au | 0.85/0.79 |
| Leaf N | cbgb.us | 0.98/0.92 |
| Leaf N | cowi.ca | 0.88/0.94 |
| Leaf N | hopl.us | 0.8/0.79 |
| Leaf N | look.us | 0.27/0.33 |
| Leaf N | mcla.us | 0.82/0.81 |
| Leaf N | sgs.us | 0.46/0.55 |
| Leaf N | sier.us | 0.64/0.76 |
| Leaf P | bnch.us | 0.16/0.24 |
| Leaf P | bogong.au | 0.61/0.6 |
| Leaf P | burrawan.au | 0.85/0.79 |
| Leaf P | cbgb.us | 0.87/0.82 |
| Leaf P | cowi.ca | 0.93/0.98 |

| **Trait name** | **site_code** | **Proportional cover of species with traits under Control and NPK** |
|---|---|---|
| Leaf P | hopl.us | 0.55/0.49 |
| Leaf P | look.us | 0.15/0.12 |
| Leaf P | mcla.us | 0.67/0.75 |
| Leaf P | sgs.us | 0.42/0.45 |
| Leaf P | sier.us | 0.65/0.63 |
| SLA | bnch.us | 0.3/0.39 |
| SLA | bogong.au | 0.2/0.37 |
| SLA | burrawan.au | 0.91/0.88 |
| SLA | cbgb.us | 0.99/0.95 |
| SLA | cowi.ca | 0.93/1 |
| SLA | hopl.us | 0.9/0.94 |
| SLA | look.us | 0.41/0.6 |
| SLA | mcla.us | 0.99/0.97 |
| SLA | sgs.us | 0.66/0.72 |
| SLA | sier.us | 0.91/0.95 |
| LES with LDMC | bnch.us | 0.66/0.73 |
| LES with LDMC | bogong.au | 0.12/0.18 |
| LES with LDMC | burrawan.au | 0.86/0.8 |
| LES with LDMC | cbgb.us | 0.94/0.92 |
| LES with LDMC | cowi.ca | 0.84/0.94 |
| LES with LDMC | hopl.us | 0.78/0.77 |
| LES with LDMC | look.us | 0.92/0.97 |
| LES with LDMC | mcla.us | 0.79/0.78 |
| LES with LDMC | sgs.us | 0.63/0.49 |
| LES with LDMC | sier.us | 0.62/0.72 |

**Table S4 Model output for the fixed effects for nutrient addition effects on plant diversity.** r2m (marginal R2): proportion of variance explained by the fixed effects in the model; r2c (conditional R2): proportion of variance explained by the fixed and random effects.

| Species included | Trait | Variable | Terms | Value | Std.error | Df | t-value | P | R2m | R2c |
|---|---|---|---|---|---|---|---|---|---|---|
| Shared | LES | CWM | (Intercept) | -0.68 | 0.32 | 29 | -2.14 | 0.04 | 0.04 | 0.94 |
| Shared | LES | CWM | trtNPK | 0.41 | 0.07 | 29 | 6.01 | 0.00 | 0.04 | 0.94 |
| Shared | LES | CWM induced by ITV | (Intercept) | -0.66 | 0.28 | 29 | -2.34 | 0.03 | 0.04 | 0.97 |
| Shared | LES | CWM induced by ITV | trtNPK | 0.39 | 0.04 | 29 | 9.58 | 0.00 | 0.04 | 0.97 |
| Shared | LES | CWM induced by replacement | (Intercept) | -0.68 | 0.34 | 29 | -2.01 | 0.05 | 0.00 | 0.97 |
| Shared | LES | CWM induced by replacement | trtNPK | 0.01 | 0.05 | 29 | 0.26 | 0.79 | 0.00 | 0.97 |
| Shared | LES | FD | (Intercept) | 0.52 | 0.06 | 29 | 9.40 | 0.00 | 0.02 | 0.69 |
| Shared | LES | FD | trtNPK | -0.06 | 0.03 | 29 | -2.04 | 0.05 | 0.02 | 0.69 |
| Shared | Leaf C | CWM | (Intercept) | 467.48 | 5.57 | 29 | 83.91 | 0.00 | 0.00 | 0.89 |
| Shared | Leaf C | CWM | trtNPK | -1.37 | 1.57 | 29 | -0.88 | 0.39 | 0.00 | 0.89 |
| Shared | Leaf C | CWM induced by ITV | (Intercept) | 467.62 | 5.62 | 29 | 83.15 | 0.00 | 0.00 | 0.90 |
| Shared | Leaf C | CWM induced by ITV | trtNPK | -1.52 | 1.54 | 29 | -0.99 | 0.33 | 0.00 | 0.90 |
| Shared | Leaf C | CWM induced by replacement | (Intercept) | 467.48 | 5.23 | 29 | 89.46 | 0.00 | 0.00 | 0.96 |
| Shared | Leaf C | CWM induced by replacement | trtNPK | 0.14 | 0.84 | 29 | 0.17 | 0.86 | 0.00 | 0.96 |
| Shared | Leaf C | FD | (Intercept) | 0.49 | 0.05 | 29 | 10.75 | 0.00 | 0.00 | 0.67 |
| Shared | Leaf C | FD | trtNPK | -0.02 | 0.03 | 29 | -0.74 | 0.47 | 0.00 | 0.67 |
| Shared | Leaf K | CWM | (Intercept) | 19.54 | 2.23 | 29 | 8.75 | 0.00 | 0.01 | 0.94 |
| Shared | Leaf K | CWM | trtNPK | 1.76 | 0.47 | 29 | 3.75 | 0.00 | 0.01 | 0.94 |
| Shared | Leaf K | CWM induced by ITV | (Intercept) | 20.12 | 2.00 | 29 | 10.04 | 0.00 | 0.01 | 0.95 |
| Shared | Leaf K | CWM induced by ITV | trtNPK | 1.18 | 0.39 | 29 | 3.02 | 0.00 | 0.01 | 0.95 |
| Shared | Leaf K | CWM induced by replacement | (Intercept) | 19.54 | 2.31 | 29 | 8.46 | 0.00 | 0.00 | 0.94 |
| Shared | Leaf K | CWM induced by replacement | trtNPK | 0.58 | 0.46 | 29 | 1.25 | 0.22 | 0.00 | 0.94 |
| Shared | Leaf K | FD | (Intercept) | 0.51 | 0.05 | 29 | 9.69 | 0.00 | 0.01 | 0.61 |

| Species included | Trait | Variable | Terms | Value | Std.error | Df | t-value | P | R2m | R2c |
|---|---|---|---|---|---|---|---|---|---|---|
| Shared | Leaf K | FD | trtNPK | -0.04 | 0.03 | 29 | -1.39 | 0.17 | 0.01 | 0.61 |
| Shared | Leaf N | CWM | (Intercept) | 25.27 | 3.15 | 29 | 8.03 | 0.00 | 0.03 | 0.93 |
| Shared | Leaf N | CWM | trtNPK | 3.64 | 0.72 | 29 | 5.10 | 0.00 | 0.03 | 0.93 |
| Shared | Leaf N | CWM induced by ITV | (Intercept) | 25.57 | 2.93 | 29 | 8.73 | 0.00 | 0.03 | 0.96 |
| Shared | Leaf N | CWM induced by ITV | trtNPK | 3.34 | 0.53 | 29 | 6.26 | 0.00 | 0.03 | 0.96 |
| Shared | Leaf N | CWM induced by replacement | (Intercept) | 25.27 | 3.09 | 29 | 8.19 | 0.00 | 0.00 | 0.97 |
| Shared | Leaf N | CWM induced by replacement | trtNPK | 0.30 | 0.44 | 29 | 0.67 | 0.51 | 0.00 | 0.97 |
| Shared | Leaf N | FD | (Intercept) | 0.50 | 0.06 | 29 | 8.91 | 0.00 | 0.01 | 0.74 |
| Shared | Leaf N | FD | trtNPK | -0.04 | 0.03 | 29 | -1.27 | 0.22 | 0.01 | 0.74 |
| Shared | Leaf P | CWM | (Intercept) | 2.28 | 0.23 | 29 | 9.97 | 0.00 | 0.06 | 0.73 |
| Shared | Leaf P | CWM | trtNPK | 0.42 | 0.11 | 29 | 3.71 | 0.00 | 0.06 | 0.73 |
| Shared | Leaf P | CWM induced by ITV | (Intercept) | 2.16 | 0.19 | 29 | 11.39 | 0.00 | 0.14 | 0.79 |
| Shared | Leaf P | CWM induced by ITV | trtNPK | 0.53 | 0.09 | 29 | 6.31 | 0.00 | 0.14 | 0.79 |
| Shared | Leaf P | CWM induced by replacement | (Intercept) | 2.28 | 0.28 | 29 | 8.26 | 0.00 | 0.00 | 0.93 |
| Shared | Leaf P | CWM induced by replacement | trtNPK | -0.12 | 0.06 | 29 | -1.83 | 0.08 | 0.00 | 0.93 |
| Shared | Leaf P | FD | (Intercept) | 0.49 | 0.05 | 29 | 10.58 | 0.00 | 0.02 | 0.72 |
| Shared | Leaf P | FD | trtNPK | -0.06 | 0.03 | 29 | -2.23 | 0.03 | 0.02 | 0.72 |
| Shared | SLA | CWM | (Intercept) | 13.64 | 2.81 | 29 | 4.85 | 0.00 | 0.01 | 0.93 |
| Shared | SLA | CWM | trtNPK | 1.65 | 0.62 | 29 | 2.67 | 0.01 | 0.01 | 0.93 |
| Shared | SLA | CWM induced by ITV | (Intercept) | 14.16 | 2.99 | 29 | 4.73 | 0.00 | 0.00 | 0.97 |
| Shared | SLA | CWM induced by ITV | trtNPK | 1.13 | 0.41 | 29 | 2.78 | 0.01 | 0.00 | 0.97 |
| Shared | SLA | CWM induced by replacement | (Intercept) | 13.64 | 2.79 | 29 | 4.89 | 0.00 | 0.00 | 0.96 |
| Shared | SLA | CWM induced by replacement | trtNPK | 0.52 | 0.49 | 29 | 1.06 | 0.30 | 0.00 | 0.96 |
| Shared | SLA | FD | (Intercept) | 0.46 | 0.05 | 29 | 10.08 | 0.00 | 0.00 | 0.54 |
| Shared | SLA | FD | trtNPK | 0.00 | 0.03 | 29 | -0.09 | 0.93 | 0.00 | 0.54 |
| All with traits | LES | CWM | (Intercept) | -0.67 | 0.31 | 29 | -2.13 | 0.04 | 0.06 | 0.94 |
| All with | LES | CWM | trtNPK | 0.49 | 0.06 | 29 | 7.55 | 0.00 | 0.06 | 0.94 |

| Species included | Trait | Variable | Terms | Value | Std.error | Df | t-value | P | R2m | R2c |
|---|---|---|---|---|---|---|---|---|---|---|
| traits | | | | | | | | | | |
| All with traits | LES | FD | (Intercept) | 0.58 | 0.06 | 29 | 10.00 | 0.00 | 0.04 | 0.72 |
| All with traits | LES | FD | trtNPK | -0.08 | 0.03 | 29 | -2.82 | 0.01 | 0.04 | 0.72 |
| All with traits | Leaf C | CWM | (Intercept) | 468.96 | 5.15 | 29 | 90.98 | 0.00 | 0.01 | 0.72 |
| All with traits | Leaf C | CWM | trtNPK | -3.21 | 2.50 | 29 | -1.28 | 0.21 | 0.01 | 0.72 |
| All with traits | Leaf C | FD | (Intercept) | 0.55 | 0.04 | 29 | 14.79 | 0.00 | 0.04 | 0.35 |
| All with traits | Leaf C | FD | trtNPK | -0.06 | 0.03 | 29 | -1.81 | 0.08 | 0.04 | 0.35 |
| All with traits | Leaf K | CWM | (Intercept) | 19.31 | 2.21 | 29 | 8.72 | 0.00 | 0.04 | 0.91 |
| All with traits | Leaf K | CWM | trtNPK | 3.04 | 0.57 | 29 | 5.30 | 0.00 | 0.04 | 0.91 |
| All with traits | Leaf K | FD | (Intercept) | 0.55 | 0.06 | 29 | 9.83 | 0.00 | 0.00 | 0.67 |
| All with traits | Leaf K | FD | trtNPK | -0.03 | 0.03 | 29 | -0.96 | 0.34 | 0.00 | 0.67 |
| All with traits | Leaf N | CWM | (Intercept) | 25.87 | 3.22 | 29 | 8.03 | 0.00 | 0.02 | 0.93 |
| All with traits | Leaf N | CWM | trtNPK | 3.18 | 0.74 | 29 | 4.27 | 0.00 | 0.02 | 0.93 |
| All with traits | Leaf N | FD | (Intercept) | 0.56 | 0.06 | 29 | 9.80 | 0.00 | 0.02 | 0.64 |
| All with traits | Leaf N | FD | trtNPK | -0.06 | 0.03 | 29 | -1.70 | 0.10 | 0.02 | 0.64 |
| All with traits | Leaf P | CWM | (Intercept) | 2.28 | 0.24 | 29 | 9.57 | 0.00 | 0.08 | 0.79 |
| All with traits | Leaf P | CWM | trtNPK | 0.48 | 0.10 | 29 | 4.77 | 0.00 | 0.08 | 0.79 |
| All with traits | Leaf P | FD | (Intercept) | 0.53 | 0.05 | 29 | 11.51 | 0.00 | 0.03 | 0.54 |
| All with traits | Leaf P | FD | trtNPK | -0.06 | 0.03 | 29 | -1.98 | 0.06 | 0.03 | 0.54 |
| All with traits | SLA | CWM | (Intercept) | 13.74 | 2.81 | 29 | 4.90 | 0.00 | 0.02 | 0.79 |
| All with traits | SLA | CWM | trtNPK | 2.67 | 1.14 | 29 | 2.35 | 0.03 | 0.02 | 0.79 |
| All with traits | SLA | FD | (Intercept) | 0.54 | 0.04 | 29 | 12.93 | 0.00 | 0.00 | 0.43 |
| All with traits | SLA | FD | trtNPK | -0.01 | 0.03 | 29 | -0.22 | 0.82 | 0.00 | 0.43 |

115

**Table S5 Model output for the fixed effects for nutrient addition effects on aboveground biomass during normal, during dry and wet, and one year after dry and wet growing season, and deviation from normal level during and one year after dry and wet growing season.** Models were specified as lme (y~trt, random=~1|site/block) for biomass under normal growing seasons and lme (y~trt, random=~1|site/block/year) for other variables. All data were log-transformed to improve normality and homogeneity of variance. r2m (marginal R2): proportion of variance explained by the fixed effects in the model; r2c (conditional R2): proportion of variance explained by the fixed and random effects.

| Variable | events | terms | Value | Std.Error | DF | t.value | p | r2m | r2c | SD (year) | SD (block) | SD (sites) |
|---|---|---|---|---|---|---|---|---|---|---|---|---|
| dev_mass | During dry | (Intercept) | 4.55 | 0.17 | 47 | 26.94 | 0.00 | 0.02 | 0.23 | 0.25 | 0.00 | 0.35 |
| dev_mass | During dry | trtNPK | 0.24 | 0.17 | 47 | 1.44 | 0.16 | 0.02 | 0.23 | 0.25 | 0.00 | 0.35 |
| dev_mass | During wet | (Intercept) | 4.54 | 0.24 | 56 | 19.23 | 0.00 | 0.03 | 0.23 | 0.00 | 0.00 | 0.57 |
| dev_mass | During wet | trtNPK | 0.47 | 0.21 | 56 | 2.27 | 0.03 | 0.03 | 0.23 | 0.00 | 0.00 | 0.57 |
| dev_mass | One year after dry | (Intercept) | 3.96 | 0.29 | 32 | 13.76 | 0.00 | 0.05 | 0.30 | 0.00 | 0.00 | 0.67 |
| dev_mass | One year after dry | trtNPK | 0.57 | 0.27 | 32 | 2.10 | 0.04 | 0.05 | 0.30 | 0.00 | 0.00 | 0.67 |
| dev_mass | One year after wet | (Intercept) | 4.42 | 0.27 | 38 | 16.19 | 0.00 | 0.05 | 0.42 | 0.00 | 0.00 | 0.73 |
| dev_mass | One year after wet | trtNPK | 0.51 | 0.20 | 38 | 2.53 | 0.02 | 0.05 | 0.42 | 0.00 | 0.00 | 0.73 |
| live_mass | During dry | (Intercept) | 5.19 | 0.27 | 47 | 19.15 | 0.00 | 0.02 | 0.75 | 0.11 | 0.19 | 0.82 |
| live_mass | During dry | trtNPK | 0.27 | 0.10 | 47 | 2.72 | 0.01 | 0.02 | 0.75 | 0.11 | 0.19 | 0.82 |
| live_mass | During wet | (Intercept) | 5.62 | 0.20 | 56 | 27.49 | 0.00 | 0.08 | 0.71 | 0.28 | 0.00 | 0.60 |
| live_mass | During wet | trtNPK | 0.49 | 0.09 | 56 | 5.69 | 0.00 | 0.08 | 0.71 | 0.28 | 0.00 | 0.60 |
| live_mass | One year after dry | (Intercept) | 5.53 | 0.18 | 32 | 30.57 | 0.00 | 0.11 | 0.74 | 0.16 | 0.00 | 0.53 |
| live_mass | One year after | trtNPK | 0.46 | 0.09 | 32 | 5.29 | 0.00 | 0.11 | 0.74 | 0.16 | 0.00 | 0.53 |

| Variable | events | terms | Value | Std.Error | DF | t.value | p | r2m | r2c | SD (year) | SD (block) | SD (sites) |
|---|---|---|---|---|---|---|---|---|---|---|---|---|
| | dry | | | | | | | | | | | |
| live_mass | One year after wet | (Intercept) | 5.68 | 0.26 | 38 | 22.01 | 0.00 | 0.05 | 0.76 | 0.13 | 0.10 | 0.77 |
| live_mass | One year after wet | trtNPK | 0.42 | 0.10 | 38 | 4.00 | 0.00 | 0.05 | 0.76 | 0.13 | 0.10 | 0.77 |
| Average mass | During normal | (Intercept) | 5.56 | 0.17 | 29 | 32.95 | 0.00 | 0.14 | 0.90 | NA | 0.13 | 0.52 |
| Average mass | During normal | trtNPK | 0.45 | 0.05 | 29 | 9.19 | 0.00 | 0.14 | 0.90 | NA | 0.13 | 0.52 |

**Table S6 Model output for the fixed effects for nutrient addition effects on facets of stability.** Models were specified as lme (y~trt, random=~1|site) or lme (y~trt*duration, random=~1|site). All stability facets were log-transformed to improve normality and homogeneity of variance. Invariability.d: detrended temporal invariability; Resistance.d/resistance.w: resistance during dry/wet growing seasons; recovery.d/ recovery.w: recovery after dry/wet growing seasons. r2m (marginal R2): proportion of variance explained by the fixed effects in the model; r2c (conditional R2): proportion of variance explained by the fixed and random effects; N sites: number of sites available for this response variable.

| Variable | Terms | Value | Std.Error | DF | t-value | p | R2m | R2c | N.sites |
|---|---|---|---|---|---|---|---|---|---|
| log.invariability.d | (Intercept) | 0.79 | 0.08 | 29 | 9.78 | 0.00 | 0.01 | 0.52 | 10 |
| log.invariability.d | trtNPK | -0.05 | 0.06 | 29 | -0.94 | 0.35 | 0.01 | 0.52 | 10 |
| log.recovery.d | (Intercept) | 0.83 | 0.25 | 29 | 3.36 | 0.00 | 0.03 | 0.08 | 10 |
| log.recovery.d | trtNPK | -0.47 | 0.33 | 29 | -1.43 | 0.16 | 0.03 | 0.08 | 10 |
| log.recovery.w | (Intercept) | 0.31 | 0.25 | 29 | 1.26 | 0.22 | 0.02 | 0.13 | 10 |
| log.recovery.w | trtNPK | -0.31 | 0.29 | 29 | -1.05 | 0.30 | 0.02 | 0.13 | 10 |
| log.resistance.d | (Intercept) | 1.04 | 0.18 | 29 | 5.65 | 0.00 | 0.02 | 0.29 | 10 |
| log.resistance.d | trtNPK | 0.25 | 0.18 | 29 | 1.39 | 0.18 | 0.02 | 0.29 | 10 |
| log.resistance.w | (Intercept) | 1.03 | 0.31 | 29 | 3.36 | 0.00 | 0.01 | 0.40 | 10 |
| log.resistance.w | trtNPK | 0.20 | 0.25 | 29 | 0.78 | 0.44 | 0.01 | 0.40 | 10 |

**Table S7 Variance explained for component models in SEMs and goodness of model fit**. Invariability.d: detrended temporal invariability; Resistance.d/resistance.w: resistance during dry/wet growing seasons; recovery.d/ recovery.w: recovery after dry/wet growing seasons. N sites: number of sites available for this response variable. r2m (marginal R2): proportion of variance explained by the fixed effects in the model; r2c (conditional R2): proportion of variance explained by the fixed and random effects.

| Source for traits | Trait name | Response | R2m | R2c | Fisher. C | df | p |
|---|---|---|---|---|---|---|---|
| Globaldatabases _genus | LES with LDMC | log.invariability.d | 0.35 | 0.56 | 15.295 | 10 | 0.122 |
| Globaldatabases _genus | LES with LDMC | log.resistance.d | 0.11 | 0.48 | 15.295 | 10 | 0.122 |
| Globaldatabases _genus | LES with LDMC | log.resistance.w | 0.20 | 0.41 | 15.295 | 10 | 0.122 |
| Globaldatabases _genus | LES with LDMC | log.recovery.d | 0.04 | 0.14 | 15.295 | 10 | 0.122 |
| Globaldatabases _genus | LES with LDMC | log.recovery.w | 0.04 | 0.19 | 15.295 | 10 | 0.122 |
| Globaldatabases _genus | LES with LDMC | richness | 0.03 | 0.88 | 15.295 | 10 | 0.122 |
| Globaldatabases _genus | LES with LDMC | CWM | 0.00 | 0.95 | 15.295 | 10 | 0.122 |
| Globaldatabases _genus | LES with LDMC | FD | 0.03 | 0.73 | 15.295 | 10 | 0.122 |
| Globaldatabases _species | Leaf N | log.invariability.d | 0.26 | 0.61 | 13.031 | 10 | 0.222 |
| Globaldatabases _species | Leaf N | log.resistance.d | 0.28 | 0.56 | 13.031 | 10 | 0.222 |
| Globaldatabases _species | Leaf N | log.resistance.w | 0.12 | 0.39 | 13.031 | 10 | 0.222 |
| Globaldatabases _species | Leaf N | log.recovery.d | 0.07 | 0.20 | 13.031 | 10 | 0.222 |
| Globaldatabases _species | Leaf N | log.recovery.w | 0.03 | 0.17 | 13.031 | 10 | 0.222 |
| Globaldatabases _species | Leaf N | richness | 0.03 | 0.88 | 13.031 | 10 | 0.222 |
| Globaldatabases _species | Leaf N | CWM | 0.00 | 0.94 | 13.031 | 10 | 0.222 |
| Globaldatabases _species | Leaf N | FD | 0.01 | 0.77 | 13.031 | 10 | 0.222 |
| Globaldatabases _species | SLA | log.invariability.d | 0.31 | 0.57 | 8.691 | 10 | 0.562 |
| Globaldatabases _species | SLA | log.resistance.d | 0.08 | 0.41 | 8.691 | 10 | 0.562 |
| Globaldatabases | SLA | log.resistance.w | 0.19 | 0.44 | 8.691 | 10 | 0.562 |

| Source for traits | Trait name | Response | R2m | R2c | Fisher. C | df | p |
|---|---|---|---|---|---|---|---|
| _species | | | | | | | |
| Globaldatabases _species | SLA | log.recovery.d | 0.04 | 0.13 | 8.691 | 10 | 0.562 |
| Globaldatabases _species | SLA | log.recovery.w | 0.03 | 0.20 | 8.691 | 10 | 0.562 |
| Globaldatabases _species | SLA | richness | 0.03 | 0.89 | 8.691 | 10 | 0.562 |
| Globaldatabases _species | SLA | CWM | 0.00 | 0.94 | 8.691 | 10 | 0.562 |
| Globaldatabases _species | SLA | FD | 0.02 | 0.72 | 8.691 | 10 | 0.562 |
| NutNet | Leaf C | log.invariability.d | 0.26 | 0.62 | 5.991 | 10 | 0.816 |
| NutNet | Leaf C | log.resistance.d | 0.11 | 0.32 | 5.991 | 10 | 0.816 |
| NutNet | Leaf C | log.resistance.w | 0.05 | 0.36 | 5.991 | 10 | 0.816 |
| NutNet | Leaf C | log.recovery.d | 0.11 | 0.27 | 5.991 | 10 | 0.816 |
| NutNet | Leaf C | log.recovery.w | 0.02 | 0.15 | 5.991 | 10 | 0.816 |
| NutNet | Leaf C | richness | 0.03 | 0.89 | 5.991 | 10 | 0.816 |
| NutNet | Leaf C | CWM | 0.01 | 0.72 | 5.991 | 10 | 0.816 |
| NutNet | Leaf C | FD | 0.04 | 0.35 | 5.991 | 10 | 0.816 |
| NutNet | Leaf K | log.invariability.d | 0.33 | 0.60 | 11.413 | 10 | 0.326 |
| NutNet | Leaf K | log.resistance.d | 0.25 | 0.49 | 11.413 | 10 | 0.326 |
| NutNet | Leaf K | log.resistance.w | 0.12 | 0.40 | 11.413 | 10 | 0.326 |
| NutNet | Leaf K | log.recovery.d | 0.07 | 0.23 | 11.413 | 10 | 0.326 |
| NutNet | Leaf K | log.recovery.w | 0.07 | 0.24 | 11.413 | 10 | 0.326 |
| NutNet | Leaf K | richness | 0.03 | 0.89 | 11.413 | 10 | 0.326 |
| NutNet | Leaf K | CWM | 0.04 | 0.91 | 11.413 | 10 | 0.326 |
| NutNet | Leaf K | FD | 0.01 | 0.67 | 11.413 | 10 | 0.326 |
| NutNet | Leaf N | log.invariability.d | 0.26 | 0.59 | 15.962 | 10 | 0.101 |
| NutNet | Leaf N | log.resistance.d | 0.09 | 0.39 | 15.962 | 10 | 0.101 |
| NutNet | Leaf N | log.resistance.w | 0.19 | 0.63 | 15.962 | 10 | 0.101 |
| NutNet | Leaf N | log.recovery.d | 0.06 | 0.08 | 15.962 | 10 | 0.101 |
| NutNet | Leaf N | log.recovery.w | 0.13 | 0.20 | 15.962 | 10 | 0.101 |
| NutNet | Leaf N | richness | 0.03 | 0.89 | 15.962 | 10 | 0.101 |
| NutNet | Leaf N | CWM | 0.02 | 0.93 | 15.962 | 10 | 0.101 |
| NutNet | Leaf N | FD | 0.02 | 0.64 | 15.962 | 10 | 0.101 |
| NutNet | Leaf P | log.invariability.d | 0.25 | 0.61 | 9.710 | 10 | 0.466 |
| NutNet | Leaf P | log.resistance.d | 0.15 | 0.41 | 9.710 | 10 | 0.466 |
| NutNet | Leaf P | log.resistance.w | 0.03 | 0.41 | 9.710 | 10 | 0.466 |
| NutNet | Leaf P | log.recovery.d | 0.24 | 0.58 | 9.710 | 10 | 0.466 |
| NutNet | Leaf P | log.recovery.w | 0.06 | 0.18 | 9.710 | 10 | 0.466 |
| NutNet | Leaf P | richness | 0.03 | 0.89 | 9.710 | 10 | 0.466 |
| NutNet | Leaf P | CWM | 0.08 | 0.79 | 9.710 | 10 | 0.466 |
| NutNet | Leaf P | FD | 0.03 | 0.54 | 9.710 | 10 | 0.466 |
| NutNet | LES | log.invariability.d | 0.28 | 0.58 | 11.121 | 10 | 0.348 |
| NutNet | LES | log.resistance.d | 0.17 | 0.49 | 11.121 | 10 | 0.348 |

| Source for traits | Trait name | Response | R2m | R2c | Fisher. C | df | p |
|---|---|---|---|---|---|---|---|
| NutNet | LES | log.resistance.w | 0.15 | 0.36 | 11.121 | 10 | 0.348 |
| NutNet | LES | log.recovery.d | 0.11 | 0.34 | 11.121 | 10 | 0.348 |
| NutNet | LES | log.recovery.w | 0.03 | 0.20 | 11.121 | 10 | 0.348 |
| NutNet | LES | richness | 0.03 | 0.89 | 11.121 | 10 | 0.348 |
| NutNet | LES | CWM | 0.06 | 0.94 | 11.121 | 10 | 0.348 |
| NutNet | LES | FD | 0.04 | 0.72 | 11.121 | 10 | 0.348 |
| NutNet | SLA | log.invariability.d | 0.23 | 0.60 | 15.834 | 10 | 0.104 |
| NutNet | SLA | log.resistance.d | 0.11 | 0.42 | 15.834 | 10 | 0.104 |
| NutNet | SLA | log.resistance.w | 0.02 | 0.40 | 15.834 | 10 | 0.104 |
| NutNet | SLA | log.recovery.d | 0.08 | 0.13 | 15.834 | 10 | 0.104 |
| NutNet | SLA | log.recovery.w | 0.02 | 0.17 | 15.834 | 10 | 0.104 |
| NutNet | SLA | richness | 0.03 | 0.89 | 15.834 | 10 | 0.104 |
| NutNet | SLA | CWM | 0.02 | 0.79 | 15.834 | 10 | 0.104 |
| NutNet | SLA | FD | 0.00 | 0.43 | 15.834 | 10 | 0.104 |

**Table S8 Partitioning direct and diversity-mediated indirect effects of nutrient addition on facets of stability based on traits extracted from global trait databases.** Results are summarized from both significant and non-significant paths in structure equation models in Fig. S5, Fig. S6, Fig. S7.

| Trait names | Effects/pathways | Resistance during dry growing seasons | Resistance during wet growing seasons | Recovery after dry growing seasons | Recovery after wet growing seasons | Temporal invariability |
|---|---|---|---|---|---|---|
| LES with LDMC | Direct effects | 0.05 | 0.15 | -0.20 | -0.12 | -0.09 |
| LES with LDMC | Direct effects through resistance and recovery | NA | NA | NA | NA | 0.02 |
| LES with LDMC | Indirect effects through diversity facets | 0.09 | -0.08 | 0.00 | 0.00 | -0.02 |
| LES with LDMC | Indirect effects through stability facets | NA | NA | NA | NA | 0.01 |
| LES with LDMC | Total effects | 0.14 | 0.08 | -0.20 | -0.11 | -0.09 |
| Leaf N | Direct effects | 0.03 | 0.13 | -0.15 | -0.12 | -0.07 |
| Leaf N | Direct effects through resistance and recovery | NA | NA | NA | NA | 0.00 |
| Leaf N | Indirect effects through diversity facets | 0.12 | -0.05 | -0.03 | -0.01 | -0.05 |
| Leaf N | Indirect effects through stability facets | NA | NA | NA | NA | 0.03 |
| Leaf N | Total effects | 0.16 | 0.08 | -0.18 | -0.13 | -0.09 |
| SLA | Direct effects | 0.11 | 0.03 | -0.15 | -0.11 | -0.06 |
| SLA | Direct effects through | NA | NA | NA | NA | 0.01 |

| Trait names | Effects/pathways | Resistance during dry growing seasons | Resistance during wet growing seasons | Recovery after dry growing seasons | Recovery after wet growing seasons | Temporal invariability |
|---|---|---|---|---|---|---|
| | resistance and recovery | | | | | |
| SLA | Indirect effects through diversity facets | 0.04 | 0.05 | -0.03 | -0.02 | -0.06 |
| SLA | Indirect effects through stability facets | NA | NA | NA | NA | 0.03 |
| SLA | Total effects | 0.16 | 0.08 | -0.18 | -0.13 | -0.09 |

**Table S9 Principal investigators contributing data but are not authors; site names match that in Table S2.** Their effort in providing data is critical to this manuscript.

| site_code | PI name | Institution |
|---|---|---|
| bogong.au | JOHN MORGAN | La Trobe University |
| burrawan.au | JENNIFER FIRN | Queensland University of Technology |
| cbgb.us | LORI BIEDERMAN | Iowa State University |
| cbgb.us | KIRSTEN HOFMOCKEL | Iowa State University |
| cbgb.us | LAUREN SULLIVAN | Iowa State University |
| sgs.us | DANA BLUMENTHAL | USDA-ARS |
| sgs.us | CYNTHIA BROWN | Colorado State University |
| sgs.us | JULIA KLEIN | Colorado State University |
| sgs.us | ALAN KNAPP | Colorado State University |

152 **Table S10 Details for contribution of each author to the manuscript.**

| Full name | Site(s) used in analysis | Developed and framed research question(s) | Analyzed data | Contributed to data analyses | Wrote the paper | Contributed to paper writing | Site coordinator | Nutrient Network coordinator | Site-level acknowledgments (funding, access, etc) |
|---|---|---|---|---|---|---|---|---|---|
| Qingqing Chen | NA | x | x | NA | x | x | NA | NA | NA |
| Yann Hautier | Frue.ch | x | x | NA | NA | x | x | NA | NA |
| Shaopeng Wang | NA | x | NA | x | NA | x | NA | NA | NA |
| Pablo Luis Peri | Potrok.ar | NA | NA | NA | NA | NA | NA | NA | NA |
| Joslin L. Moore | bogong.au | NA | NA | NA | NA | x | x | NA | NA |
| Andrew S. MacDougall | cowichan | NA | NA | NA | NA | x | x | NA | NA |
| Sally Power | NA | NA | NA | NA | NA | x | x | NA | NA |
| Jonathan D. Bakker | smith.us | NA | NA | NA | NA | x | x | NA | NA |
| Anne Ebeling | jena.de | NA | NA | NA | NA | x | x | NA | NA |
| Christiane Roscher | jena.de | NA | NA | NA | NA | x | x | NA | NA |
| Maria C Caldei | comp.pt | NA | NA | NA | NA | x | x | NA | Portuguese Science Foundation |

| Full name | Site(s) used in analysis | Developed and framed research question(s) | Analyzed data | Contributed to data analyses | Wrote the paper | Contributed to paper writing | Site coordinator | Nutrient Network coordinator | Site-level acknowledgments (funding, access, etc) |
|---|---|---|---|---|---|---|---|---|---|
| ra | | | | | | | | | (FCT) for funding the research unit CEF (UIDB/00239/2020) and to Companhia das Lezirias for field access |
| Carla Nogueira | comp.pt | NA | NA | NA | NA | x | x | NA | NA |
| Eric W. Seabloom | bnch.us, hopl.us, look.us, mcla.us, sier.us | NA | NA | NA | NA | x | x | x | NA |
| Carly J. Stevens | lancaster.uk | NA | NA | NA | NA | x | x | NA | NA |
| Sumanta Bagchi | NA | NA | NA | x | NA | x | x | NA | I can help you revise/upgrade the SEM in order to present a test of alternative competing hypotheses in Fig4. Current ms has one hypothesis, but it is not clear whether the data also fit an alternative "null" |

| Full name | Site(s) used in analysis | Developed and framed research question(s) | Analyzed data | Contributed to data analyses | Wrote the paper | Contributed to paper writing | Site coordinator | Nutrient Network coordinator | Site-level acknowledgments (funding, access, etc) |
|---|---|---|---|---|---|---|---|---|---|
| | | | | | | | | | model. This can be evaluated with competing SEMs where Fig1 is "not true". |
| Elizabeth T. Borer | bnch.us, hopl.us, look.us, mcla.us, sier.us | NA | NA | NA | NA | x | x | x | NA |
| Forest Isbell | NA | NA | NA | x | NA | x | NA | NA | NA |
| Ian Donohue | NA | NA | NA | NA | NA | x | NA | NA | NA |
| Juan Alberti | NA | NA | NA | x | NA | x | NA | NA | NA |
| Anke Jentsch | NA | NA | NA | NA | NA | x | x | NA | Federal Ministry of Education and Research (BMBF, SUSALPS; FKZ 031B0516C) and Upper Franconian Trust (Oberfrankenstiftung; grant number: OFS FP00237). |
| Michelle Tedder | gilb.za | NA | NA | NA | NA | x | x | NA | NA |
| Kevin Kirkman | gilb.za | NA | NA | NA | NA | x | x | NA | NA |

| Full name | Site(s) used in analysis | Developed and framed research question(s) | Analyzed data | Contributed to data analyses | Wrote the paper | Contributed to paper writing | Site coordinator | Nutrient Network coordinator | Site-level acknowledgments (funding, access, etc) |
|---|---|---|---|---|---|---|---|---|---|
| Siddharth Bharath | NA | NA | NA | NA | NA | NA | NA | NA | NA |
